# 3D models of Alzheimer’s disease patient microglia recapitulate disease phenotype and show differential drug responses compared to 2D

**DOI:** 10.1101/2021.03.17.435758

**Authors:** Carla Cuní-López, Hazel Quek, Lotta E. Oikari, Romal Stewart, Tam Hong Nguyen, Yifan Sun, Christine C. Guo, Michelle K. Lupton, Anthony R. White

## Abstract

Alzheimer’s disease (AD) is an incurable neurodegenerative disorder with a rapidly increasing prevalence worldwide. Current approaches targeting hallmark pathological features of AD have had no consistent clinical benefit. Neuroinflammation is a major contributor to neurodegeneration and hence, microglia, the brain’s resident immune cells, are an attractive target for potentially more effective therapeutic strategies. However, there is no current *in vitro* model system that faithfully recapitulates patient-specific microglial characteristics. To address this shortcoming, we developed novel 3D models of monocyte-derived microglia-like cells (MDMi) from AD patients. MDMi in 3D exhibited mature microglial features, including a highly branched morphology and enhanced bonafide microglial marker expression compared to 2D. Moreover, AD MDMi in 3D co-cultures with neuro-glial cells showed altered cell-to-cell interactions, growth factor and cytokine secretion profiles and responses to amyloid-β. Drug screening assays in 3D AD MDMi revealed different cytokine responses compared to 2D. Our study demonstrates disease- and drug-specific responses in 3D MDMi models that are not apparent in 2D and presents a new 3D platform for more effective and personalised drug testing.

## Introduction

Alzheimer’s disease (AD) is a complex age-related neurodegenerative disorder involving progressive impairment of cognitive functions, with distinct pathogenicity and clinical phenotypes among patients. Predictions estimate that AD, together with other neurodegenerative diseases, will become the second leading cause of death in the next 20 years (*1*). Nevertheless, no prevention strategies or cure exist despite major advances in deciphering the molecular basis of AD. The most characteristic neuropathological hallmarks of AD brains are the presence of extracellular senile plaques, consisting primarily of misfolded amyloid-β (Aβ) protein, and intracellular neurofibrillary tangles, composed of hyperphosphorylated tau protein. Accumulation of protein aggregates in the brain parenchyma triggers multiple deleterious processes, including oxidative stress and mitochondrial dysfunction (*2*), which alter brain homeostasis. For decades, reducing the protein aggregate burden in AD brains has been the main goal of candidate therapeutics, but these strategies have yielded poor outcomes in clinical trials. The field is therefore in desperate need for more effective drug targets.

Chronic neuroinflammation and sustained activation of pro-inflammatory pathways are critical components of many neurodegenerative diseases, including AD. Recent reports have shown that neuroinflammation plays a major role in the pathogenesis and progression of AD (*3, 4*). Microglia, the resident immune modulators of the brain, are key effectors of neuroinflammation and hence represent a promising candidate for targeted therapeutics for AD and other neurodegenerative diseases. Microglia develop aberrant phenotypes under non-homeostatic brain conditions and consequently mediate multiple pathogenic mechanisms, including neuron and synapse degeneration, that are critical to the cognitive decline characteristic of AD (*5, 6*). Moreover, most AD risk gene variants (for example *TREM2*, *APOE*, *CLU, CD33*, *PILRB*, *BIN1*, *PLCG2* and *MEF2C*) converge on biological pathways linked to microglial function (*7–12*). Recent studies have correlated the responses of diseased microglia in AD brains with the varied clinical presentations seen among patients. Indeed, the individual’s genetic makeup determines whether diseased microglia will prevent or exacerbate the pathogenic processes underlying disease in that particular patient’s brain (*13–15*). As such, microglia not only contribute to the pathology of AD but also show patient-specific characteristics, thus being essential players in the patient heterogeneity observed in AD.

Current *in vitro* model systems used to study the role of microglia in AD (reviewed in (*16, 17*)) lack either clinical relevance or physiological complexity, thereby affecting translatability of drug outcomes into the clinic. Murine microglia lack the ability to fully recapitulate disease phenotypes of AD patients due to the little resemblance of immune functions and ageing processes between mice and humans (*18–21*). Human immortalised microglia cell lines are genetically and functionally very different from *in vivo* microglia (*22–24*). Freshly isolated primary microglia from AD patients are normally obtained from post-mortem brains in low yields and rapidly lose microglial phenotypic signatures upon removal from the brain environment (*22*). Lastly, human induced pluripotent stem cell (hiPSC)-derived microglia allow for the generation of a clinically relevant, patient-specific microglia platform. However, establishing hiPSC-derived microglia requires costly, long and technically challenging protocols that result in variable differentiation efficiencies (*25*), and the cells lose patient-specific traits, including ageing markers, upon reprogramming (*26*).

The monocyte-derived microglia-like cell (MDMi) model system addresses the shortcoming of the above models and provides a novel, cost-effective approach for the rapid generation of personalised microglia cultures from living patients. This method has been previously applied by us and others using *ex vivo* blood-derived monocytes from schizophrenia (*27, 28*), Nasu-Hakola disease (*29*) and amyotrophic lateral sclerosis (ALS) (*30*) patients, demonstrating disease-associated phenotypes in the patient-derived MDMi. In addition to their controlled genetic background, MDMi are readily available and yield mature microglia in a short time frame, thus allowing for the study of mature microglia from large patient cohorts (*31, 32*).

Microglial identity is driven by the multicellular milieu and three-dimensional (3D) network of macromolecules present in the brain. Therefore, the traditional two dimensional (2D) culture conditions of microglia *in vitro* systems greatly abrogate their ability to replicate mature microglial function (*33*). The physiological relevance of the MDMi model can be increased by using 3D *in vitro* culture techniques and co-cultures with neuro-glial cells to incorporate the cues supporting microglial development *in vivo*. The development of a complete AD pathological cascade in 3D, but not in 2D, shows an improved *in vitro* disease modelling capacity of 3D culture systems compared to 2D (*34, 35*). However, no 3D *in vitro* model of AD has yet incorporated patient-derived microglial cells in a highly reproducible and experimentally flexible 3D cell culture system (*36*).

In this study, we generated for the first time 3D patient-specific MDMi models from multiple living AD patients. These 3D hydrogel-based MDMi models are consistent and easy to generate and allow for the establishment of 3D MDMi co-cultures with human neuro-glial cells, which provide a more complex and physiologically relevant culture environment. We characterised 3D MDMi models at different levels. Firstly, we examined whether a 3D hydrogel scaffold enhances microglia-like features (*i.e.,* morphology and marker expression) in MDMi compared to a 2D platform using cells from healthy controls. Secondly, we demonstrated the feasibility to generate 3D MDMi models in 3D co-culture. Thirdly, we studied AD-specific changes in the 3D cultures, including morphology, expression of AD risk genes, cell-to-cell interaction with neuro-glial cells and functional responses against Aβ aggregates. Finally, to test the potential applicability of 3D MDMi platforms in a drug screening setting, we compared drug responses in MDMi between the 2D, 3D and 3D co-culture models. Together, the utility of the 3D MDMi models presented here opens new avenues for more predictable and personalised *in vitro* microglia models to test candidate therapeutics.

## Methods

### Study cohort

This study involved the recruitment of Alzheimer’s disease (AD) and Healthy control (HC) participants through the *Prospective Imaging Studying of Aging: Genes, Brain and Behaviour study* (PISA) at QIMR Berghofer Medical Research Institute, Queensland, Australia (*37*). All research adhered to the ethical guidelines on human research outlined by the National Health and Medical Research Council of Australia (NHMRC). Ethical approval was obtained from QIMR Berghofer Medical Research Institute. All participants provided informed consent before participating in the study. The number of samples varied in each assay due to the limited proliferative capacity of MDMi in culture, and the quantity of blood samples available from each donor. Further, repeated longitudinal sampling of peripheral blood from patients was not within the scope of this study. All samples used for assays were randomly selected, with matching age, gender and apolipoprotein E (APOE) status for each assay. APOE genotyping was performed in the Sample Processing Facility at QIMR Berghofer Medical Research Institute, Queensland, Australia, as previously described (*37*).

### Isolation of peripheral blood mononuclear cells (PBMCs)

Peripheral venous blood samples were collected in ethylenediaminetetraacetic acid (EDTA) tubes (Becton-Dickson, NJ, USA). PBMCs separation was performed within 2 h of blood withdrawal using SepMate™ tubes (StemCell Technologies, BC, Canada) as per manufacturer’s instructions. PBMCs were washed twice with PBS containing 1mM EDTA and subsequently frozen in 10% dimethyl sulphoxide (DMSO) (Merck KGaA, Hesse, Germany) and 90% foetal bovine serum (ThermoFisher Scientific, CA, USA) (v/v).

### Establishment of 2D and 3D MDMi cultures

MDMi in 2D were generated as described previously (*30*). Briefly, cryopreserved PBMCs were thawed and seeded onto plates coated with Matrigel (Corning, NY, USA). After 24 h incubation at standard humidified culture conditions (37°C, 5% CO_2_), non-adherent cells were removed and a cell population enriched in monocytes remained adhered to the culture vessel. To induce MDMi differentiation, monocytes were then cultured in serum-free RPMI-1640 GlutaMAX medium (Life Technologies, Grand Island, NY, USA) supplemented with 0.1 μg/ml of interleukin (IL)-34 (IL-34) (Lonza, Basel-Stadt, Switzerland), 0.01 μg/ml of granulocyte-macrophage colony-stimulating factor (GM-CSF) (Lonza, Basel-Stadt, Switzerland) and 1% (v/v) penicillin/streptomycin (Life Technologies, Grand Island, NY, USA) for 14 days.

To induce MDMi differentiation in 3D, monocytes were resuspended in Matrigel diluted with ice-cold culture medium at a 1:3 ratio. Matrigel-cell mixtures were seeded in 96-well plates with medium containing 0.1 μg/ml IL-34 and 0.01 μg/ml GM-CSF. 3D MDMi were collected or used for downstream assays after an average of 35 days in culture.

### Establishment of 2D and 3D human neural progenitor cell (NPCs) cultures

The human ReNcell VM immortalised neural progenitor cell line (EMD Millipore, Billerica, MA, USA) was cultured as per manufacturer’s instructions, with some modifications. Briefly, cells were plated onto Matrigel-coated plates for 2D cultures or mixed with a 1:3 Matrigel dilution to initiate the 3D cultures. Cells were maintained in DMEM/F12 GlutaMAX medium (Life Technologies, Grand Island, NY, USA) containing 2% (v/v) B27 supplement, 20 μg/ml epithelial growth factor (EGF) (Sigma-Aldrich, MO, USA), 20 μg/ml fibroblast growth factor 2 (FGF-2) (Lonza, Basel-Stadt, Switzerland) and 1% (v/v) penicillin/streptomycin. Both 2D and 3D cultures were spontaneously differentiated for 1, 14 or 30 days by withdrawing growth factors from the maintenance medium (ReN base medium). All cells used were in passages 7-10 to ensure consistent spontaneous neuro-glial differentiation across independent experiments.

### Establishment of 3D co-cultures (MDMi and ReNcell VM)

ReNcell VM were plated in 3D as described above and cultured for 1 day in ReN base medium to induce spontaneous differentiation. Monocytes were embedded in a 1:3 Matrigel dilution and seeded with 3D ReNcell VM cultures at 1:2.5 to 1:5 monocyte to ReNcell VM ratios. 3D co-cultures were maintained in 50% (v/v) ReN base medium and MDMi culture medium for an average of 35 days.

### Immunocytochemistry

Immunofluorescence staining of 2D cultures was performed as described previously (*30*). MDMi and ReNcell VM were cultured on 8-well chamber slides (Ibidi, DKSH, Germany) and 13-mm plastic coverslips (Sarstedt, Nümbrecht, Germany), respectively. Cells were fixed in 4% paraformaldehyde (PFA) or ice-cold methanol for 15 min and then washed with PBS. Blocking was performed at RT with 5% bovine serum albumin (BSA) (Sigma-Aldrich, MO, USA) in PBS. Primary antibodies for TREM 2 (1:500; Abcam, # ab201621), P2RY12 (1:200; Alomone Labs, #APR-20), TMEM119 (1:400; Abcam, # ab185333), IBA1 (1:500; Wako, #019-19741), Nestin (1:200; Abcam, #ab22035), GFAP (1:2000; Abcam, #ab4674), GalC (1:50; Santa Cruz, #sc-518055), Doublecortin (DCX) (1:200; Abcam, ab36447) and βIII-tubulin (TUBB3) (1:500; BioLegend, #801202) were diluted in blocking solution and incubated overnight at 4°C. Cells were then washed three times with 0.1% Triton-X 100 in PBS and incubated with 1:250 Alexa Fluor 488 (#A-11034) / 594 (#A-21203) / 647 (#A-21244) secondary antibodies (ThermoFisher Scientific, CA, USA) for 2 h at RT in the dark and counterstained with a nuclear dye (Hoechst 33342, 1 μg/ml). Immunofluorescence staining of 3D cultures was performed according to (*38*) with some modifications. 3D cultures, established on black optical 96-well plates (ThermoFisher Scientific, CA, USA), were fixed with 4% PFA overnight at RT and washed twice with PBS. Cultures were then permeabilised for 30 min with 0.3% Triton-X 100 in PBS, rinsed with PBS and blocked overnight at 4°C with 2% BSA (Sigma-Aldrich, MO, USA) in PBS. Primary antibody solutions were added as described above and incubated for 24 h at 4°C. Secondary antibody solutions were incubated for 5 h at RT in the dark. Following primary and secondary antibody incubations, cultures were washed five times (10 min each) with 0.1% Triton-X 100 in PBS and counterstained with Hoechst 33342. Images were captured using a confocal laser scanning microscope (LSM-780, Carl Zeiss) at 20X and 40X magnification and processed using the Zeiss ZEN software.

### RNA extraction and quantitative real-time PCR (qRT-PCR)

RNA and cDNA were prepared as previously described (*39*). Total RNA was extracted using a Direct-zol RNA Miniprep kit (Integrated Sciences, Australia) as per manufacturer’s protocol. Conversion to cDNA was carried out using a SensiFAST™ cDNA synthesis kit (Bioline, London, UK). For qRT-PCR, cDNA was diluted 1:10 to generate working solutions and combined with SensiFAST™ SYBR® Lo-ROX master mix and gene-specific primers (see primer sequences in **Table S1**). The qRT-PCR runs were performed as triplicate on Applied Biosystems ViiA 7 (ThermoFisher Scientific, CA, USA). Endogenous control *18S* was used as a housekeeping gene for normalisation. Relative gene expression levels were calculated using the ΔΔCt method.

**Table 1.** Summary of donor information.

| Study cohorts |  | Healthy Control (HC) | Alzheimer's disease (AD) |
| --- | --- | --- | --- |
| N° of participants |  | *n* = 12 | *n* = 13 |
| Sex of participants | Females (%) | 58.3% (7/12) | 53.8% (7/13) |
|  | Males (%) | 41.7% (5/12) | 46.2% (6/13) |
| Age of participants (mean ± SD) |  | 68.5 ± 2.7 | 69.3 ± 5.4 |
| APOE genotype | E3/E3 (%) | 25% (3/12) | 7.7% (1/13) |
|  | E3/E4 (%) | 58.3% (7/12) | 69.2% (9/13) |
|  | E4/E4 (%) | 16.7% (2/12) | 23.1% (3/13) |

SD = standard deviation
APOE = apolipoprotein E

### Multiplex bead-based immunoassay

The LEGENDplex^TM^ Human Inflammation (#740809) and Growth Factor (#740180) kits (BioLegend, CA, USA) were used to detect cytokines and growth factors in conditioned media from 3D MDMi and ReNcell VM mono- and co-cultures. The assay was performed as per manufacturer’s instructions. Briefly, conditioned media were incubated with a cocktail of antibody-conjugated capture beads. Then biotinylated detection antibodies were added followed by streptavidin-phycoerythrin (SA-PE). The amount of analytes of interest in the samples was calculated as a proportion of the fluorescent signal intensity provided by capture bead-analyte-detection antibody-SA-PE sandwiches. Signals were acquired on a BD LSRFortessa 5 (BD Biosciences, CA, USA) using FACSDiva software, and analysed using Qognit, a cloud-based LEGENDplex^TM^ software (BioLegend, CA, USA). Concentrations (pg/ml) were normalised to total amount of protein in the cultures.

### Morphology analysis

Quantification of morphological parameters of MDMi in 2D and 3D mono-cultures was performed by adapting a previous method (*40*). In brief, phase contrast images acquired with a 20X objective were processed in FIJI software (National Institutes of Health, Maryland, USA) using a macro script that applied a threshold, followed by processing functions “despeckle”, “close” and “remove outliers” that generated a binary image. Binary images were then run on the *AnalyseSkeleton(2D/3D)* plugin, which resulted in skeletonised images. The “results and branch information” outputs from the plugin contained data on branch length, endpoint number and triple and quadruple junctions number. Binary images were also analysed using the *Analyze particles* function in FIJI. This calculated the “solidity” or “ramification index” value, which results from dividing the area of MDMi by its convex area (*i.e.,* area of the smallest polygon drawn around the cell). More ramified cells have a bigger convex area and thus a smaller ramification index (<1). Mean single cell values for each parameter were calculated. The total number of MDMi analysed per donor was 100 in 2D and 20 in 3D.

### Cell contacts analysis in 3D co-cultures

Confocal Z-stack images of 3D co-cultures acquired with a 20X objective were rendered in 3D using the Imaris software (Bitplane, Belfast, UK). During image acquisition, the Z-interval was set at “Optimal” so that the number of acquired slices was suitable for the given stack size, objective lens, and pinhole diameter. Following surface modelling using the Surface function in Imaris, the *Surface-Surface contact area* extension module was applied to measure the areas in contact between ReNcell VM and MDMi as well as the number of contacts established. Both parameters were then normalised to the total number of MDMi in the image. A total of 200 MDMi in co-culture were analysed for each donor in the HC and AD cohorts.

### Preparation of amyloid-β (Aβ) fibrils

Fluorescein isothiocyanate (FITC)-conjugated amyloid-β peptides 1-42 (FITC-Aβ_1-42_) (Bachem, M2585, CH) were dissolved in DMSO to a concentration of 500μM and stored at −80°C. FITC-Aβ_1-42_ were incubated for 24 h at 37°C in the 3D cultures prior to imaging to allow for the fibrillisation of the peptides and formation of Aβ fibrillary aggregates.

### Aβ aggregates exposure and surveillance analysis

FITC-Aβ_1-42_ were added at 5μM to MDMi 3D mono- and co-cultures at day 35 of differentiation. After 24 h, cultures were imaged on an EVOS FL Auto 2 (ThermoFisher Scientific, CA, USA). Scans were set to image multiple z-stack planes every 12 h for 7 days using a 10X objective. At least 3 fields of view were scanned per well. MDMi located within an area of 90,000 μm^2^ containing one or more Aβ fibrillary aggregate were tracked using the *Manual tracking* plugin in FIJI. Migrated distance and velocity of tracked cells were calculated and normalised to the number of MDMi in the analysed areas. Between 100-200 MDMi per individual was tracked in both HC and AD cohorts.

### Statistical analysis

All statistical analyses were performed using GraphPad Prism software version 8 (Graphpad Software, CA, USA). Comparisons between two groups were analysed with two-tailed Student’s *t*-test or Mann-Whitney *U*, when normality assumptions were not met. Comparisons between three or more groups were analysed by one- or two-way analysis of variance (ANOVA) followed by post-hoc tests. Data are presented as mean ± SEM or mean ± SD and *P* ≤ 0.05 was considered significant. Statistical significance was determined as \**P* < 0.05, \*\**P* < 0.01, \*\*\**P* < 0.001, \*\*\*\**P* < 0.0001, as detailed in figure legends.

## Results

### MDMi in 3D show increased survival and more mature microglial features compared to 2D

We have previously differentiated monocytes into MDMi in a 2D platform using Matrigel-coated plates (*30*). To develop a more physiologically relevant MDMi model with a better representation of the 3D structure of the brain, we differentiated monocytes into MDMi in a 3D platform. We embedded the monocytes in Matrigel (**Fig. 1A**; **Fig. S1A**), resulting in cultures with 6.2-fold higher cell thickness (*i.e.,* size of the Z-stack that captured the whole cell) compared to 2D (**Fig. 1B**; **Fig. S1B; Movie S1, 2**). Remarkably, MDMi survival in 3D was significantly increased by 2.5 fold compared to 2D (**Fig. 1C**). Hence, MDMi were cultured for an average of 14 days in 2D and 35 days in 3D, after which features of cellular ageing, including enlarged cell sizes and increased vacuolisation, were observed. We next examined if the 3D culture conditions affected microglial features in MDMi, including morphology and expression of microglia-enriched markers, compared to 2D. Overall, we observed that 3D MDMi showed a highly ramified branched structure and increased branch complexity compared to 2D (**Fig. 1D**). Branched structure parameters, such as branch length and number of branches (endpoints), and branch complexity parameters, such as number of triple and quadruple junctions (points at which branches divide into three or four sub-branches, respectively), were significantly increased in 3D compared to 2D MDMi (**Fig. 1E-G**). Ramified microglia have a highly branched morphology with a larger convex area than ameboid cells, thus correlating with a low ramification index (*41*). We observed that the ramification index of 3D MDMi was lower than 2D MDMi (**Fig. 1H**), confirming an enhanced ramified morphology of MDMi in the 3D platform. Such enhancement in MDMi ramified morphology could be likely due to the larger surface area for growth and differentiation provided by the 3D Matrigel scaffold.

**Fig. 1.**
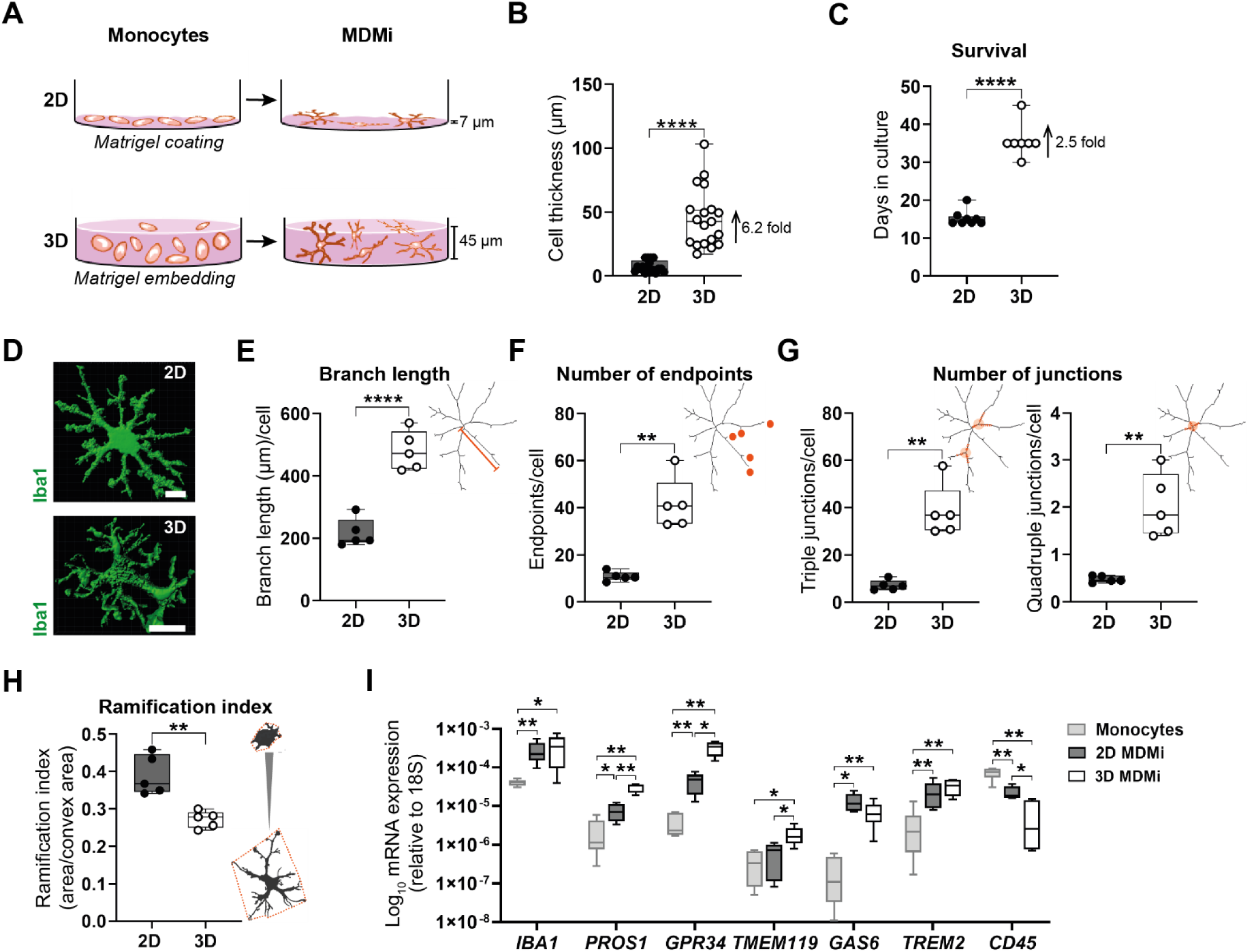
Generation and characterisation of distinctive microglia features in 2D and 3D MDMi. **(A)** Schematic illustration of monocyte differentiation into MDMi in 2D, achieved by seeding the monocytes on a Matrigel-coated surface, or in 3D, achieved by embedding the monocytes in a thick Matrigel layer. **(B)** Cell thickness of MDMi in 2D (*n* = 17 independent cultures) and 3D (*n* = 19 independent cultures) cultures. **(C)** Survival of 2D (*n* = 8) and 3D MDMi (*n* = 7). **(D)** 3D surface rendered images of 2D and 3D MDMi stained for Iba1 (green). Scale bars, 25 μm. Quantification of morphological parameters in 2D (*n* = 5) and 3D MDMi (*n* = 5), including **(E)** branch length, **(F)** number of endpoints, **(G)** number of junctions, including triple junctions (left) and quadruple junctions (right) and **(H)** ramification index (area/convex area). Representative skeleton and binary images are included on the right of each graph to illustrate the morphological measurements. **(I)** mRNA expression of microglia-(*IBA1*, *PROS1, GPR34*, *TMEM119*, *GAS6*, *TREM2*) and leukocyte-enriched (*CD45*) markers in the starting monocyte population (*n* = 6) and the resulting 2D (*n* = 7) and 3D (*n* = MDMi cultures. Data are presented as mean ± SD. Each single data point represents one biological replicate. Unpaired Student’s *t* test with or without Welch’s correction, two-tailed; one-way ANOVA with Tukey’s multiple comparison test in **I**; \**P* < 0.05, \*\**P* < 0.01, \*\*\*\**P* < 0.0001.

We have previously reported that MDMi cultured in 2D showed a microglial phenotype compared to monocytes demonstrated by the upregulated expression of microglia-enriched markers, including *PROS1*, *GPR34*, *GAS6* and *TREM2*, and the downregulated expression of the leukocyte marker *CD45* (*30*). Expectedly, all seminal microglial markers (*IBA1, PROS1*, *GPR34*, *TMEM119*, *GAS6* and *TREM2*) were upregulated and *CD45* was downregulated in 3D MDMi compared to monocytes (**Fig. 1I**). Interestingly, in comparison to 2D, we observed a significantly increased expression of *PROS1*, *GPR34* and *TMEM119* in 3D MDMi, while similar expression levels were observed for *IBA1*, *GAS6* and *TREM2* between 3D MDMi and 2D MDMi (**Fig. 1I**). These results suggest that the 3D culture was able to promote selective microglia-enriched markers in MDMi. Importantly, as immature microglia have been reported to lack *TMEM119* expression (*42*), the upregulation of *TMEM119* in 3D MDMi compared to 2D MDMi further indicates an enhanced microglial maturity in the 3D platform. Finally, positive immunostaining of Trem2 and P2ry12 proteins in 3D MDMi confirmed the retention of microglial proteins in MDMi for up to 35 days (**Fig. S1C**).

Overall, these results demonstrate that MDMi cultured in 3D survive longer in culture and better recapitulate microglial features, including a more ramified morphology and enhanced expression of microglial core genes, compared to 2D.

### Human neural progenitor cells differentiated in 3D generate more mature neuro-glial populations compared to 2D

To mimic the neuro-glial cues present in the brain microenvironment, we used the immortalised human neural progenitor cell (NPC) line ReNcell VM to establish a co-culture platform with MDMi. ReNcell VM are derived from the ventral mesencephalon region of a foetal brain (*43*) and give rise to mixed populations of astrocytes, oligodendrocytes and neurons upon differentiation (*43, 44*). Using immunofluorescence, we confirmed the presence of cells expressing nestin (NPC marker), GFAP (immature and mature astrocyte marker), GalC (oligodendrocyte progenitor cells (OPCs) and mature oligodendrocyte marker) and doublecortin (DCX, immature neuron marker) after 1 and 30 days of ReNcell VM differentiation in 2D (**Fig. 2A**). As ReNcell VM differentiate, they lose their stemness and proliferative capacity. This was confirmed by a significant decrease in *Ki67* expression, a proliferation marker, by day 14 of differentiation that was further decreased by day 30 of differentiation (**Fig. 2B**). Consistently, the reduction of stemness was reflected by the decreased expression of well-known markers of radial glial progenitor cells (*SOX2*, *NESTIN*, *GLAST* and *BLBP*) by day 30. In contrast, immature astrocyte, OPCs and neuron markers (*GFAP*, *PLP-1* and *TBR2*, respectively) showed increased expression trends after day 1 of differentiation. Markers indicating enhanced astrocytic (*GLT-1*), oligodendrocytic (*GalC*) and neuronal (*MAPT*, *synaptophysin* (*SYP*)) maturity showed a similar upregulation by day 30 but reached lower expression levels than those from radial glia and immature glial markers (**Fig. 2C**). Overall, these results demonstrate that differentiation of ReNcell VM generates mixed neuro-glial cell populations with predominance of glial cell types including radial glial progenitor cells, astrocytes and OPCs.

**Fig. 2.**
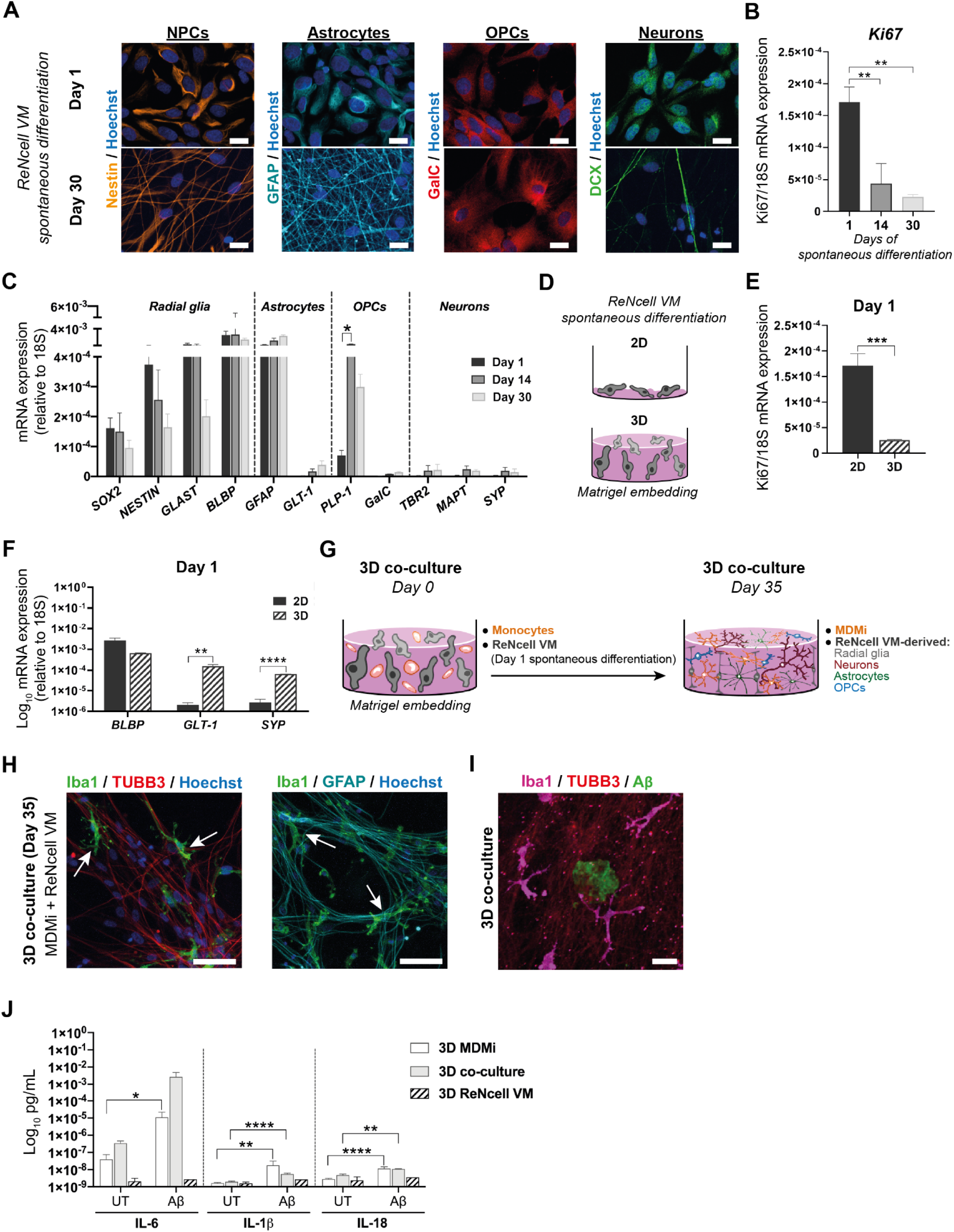
MDMi in 3D co-culture with ReNcell VM-derived neuro-glial cells exhibit inflammatory responses towards Aβ aggregates. **(A)** Immunostaining of 2D ReNcell VM cultures differentiated for 1 and 30 days shows expression of characteristic markers of neural progenitor cells (NPCs; Nestin), astrocytes (GFAP), oligodendrocyte progenitor cells (OPCs; GalC) and neurons (Doublecortin, DCX). Scale bars, 100 μm. **(B)** *Ki67* mRNA expression in 2D ReNcell VM cultures differentiated for 1, 14 or 30 days (*n* = 3 independent experiments). **(C)** mRNA expression of a panel of radial glia (NPCs), astrocytes, OPCs and neuron markers in 2D ReNcell VM at days 1, 14 and 30 of spontaneous differentiation (*n* = 3 independent experiments). **(D)** Schematic of ReNcell VM cultures undergoing spontaneous differentiation for 1 day in 2D, or in 3D upon embedment in Matrigel. mRNA expression of **(E)** the proliferation marker *Ki67* alongside the **(F)** the radial glia marker *BLBP*, the mature astrocyte marker *GLT-1* and the mature neuron marker *Synaptophysin* (*SYP*) in 2D (*n* = 3 independent experiments) and 3D (*n* = 3 independent experiments) ReNcell VM cultures spontaneously differentiated for 1 day. **(G)** Schematic depicting how the 3D co-culture is generated: monocytes pre-mixed with Matrigel are added to a 3D culture of ReNcell VM differentiated for 1 day (left). The co-culture is left to differentiate for 35 days, giving rise to a mixed population of MDMi with ReNcell VM-derived neuro-glial cells (right). **(H)** Immunofluorescence analysis of MDMi and ReNcell VM 3D co-culture at day 35 of differentiation. MDMi were stained for Iba1 (arrows) and ReNcell VM were stained for β3-tubulin (TUBB3) and GFAP. Scale bars, 100 μm. **(I)** Immunofluorescence image of 3D co-cultures containing FITC-Aβ aggregates. Scale bar, 100 μm. **(J)** Concentration of secreted pro-inflammatory cytokines IL-6, IL-1β and IL-18 by 3D MDMi (*n* = 2) and ReNcell VM (*n* = 1) mono-cultures and 3D co-cultures (*n* = 3) upon exposure to FITC-Aβ aggregates. Data are presented as mean ± SEM. One-way ANOVA with Tukey’s multiple comparison test in **B**, **C**; unpaired Student’s *t* test with or without Welch’s correction, two-tailed in **E**, **F**, **J**; \**P* < 0.05, \*\**P* < 0.01, \*\*\**P* < 0.001, \*\*\*\**P* < 0.0001.

We next examined whether differentiation under 3D Matrigel culture conditions would enhance the maturity of ReNcell VM-derived neuro-glial cell populations compared to 2D. Hence, we differentiated ReNcell VM in 2D and 3D for 1 day (**Fig. 2D**). We observed that 3D ReNcell VM showed a significant reduction of *Ki67* expression and a significant increase of mature astrocyte and neuron marker (*GLT-1* and *SYP*, respectively) expression compared to 2D (**Fig. 2E-F**). These results indicate that the 3D platform enhances the maturation of the astrocytic and neuronal populations in ReNcell VM within 1 day of differentiation compared to 2D.

Together, ReNcell VM-derived neuro-glial populations provide a multicellular brain-like environment that allows for the generation of co-cultures with MDMi, which represent more complex and physiologically relevant MDMi models.

### MDMi co-cultured in 3D with human neural progenitor cells elicit an inflammatory response to aggregated amyloid-β (Aβ)

We next generated co-cultures of MDMi and ReNcell VM. Monocytes, pre-mixed with Matrigel, were added to 3D ReNcell VM differentiated for 1 day. Both cell types were then left to differentiate together for 35 days, resulting in a mixed 3D co-culture containing MDMi and ReNcell VM-derived radial glial progenitor cells, neurons, astrocytes and OPCs (**Fig. 2G**). Interestingly, we observed that in 3D co-culture, monocytes readily differentiated into MDMi with a microglia-like morphology (**Fig. 2H**). This is in contrast to 2D co-culture, where monocytes retained their round morphology (**Fig. S2**). This indicates that the 3D platform is able to provide culture conditions suitable for co-culturing MDMi and ReNcell VM, likely due to the support of the 3D Matrigel scaffold.

Aβ aggregation is a major histopathological hallmark of AD brains. Hence, we incorporated FITC-Aβ peptides into the 3D co-cultures. Remarkably, we observed that FITC-Aβ peptides readily formed substantial aggregates within 24 h of incubation in 3D (**Fig. 2I**; **Fig. S3**) as opposed to 2D, where FITC-Aβ remained in peptides (**Fig. S3**). This suggests that 3D culture conditions may increase Aβ aggregation (**Fig. S3**), which is consistent with previous reports showing an accelerated Aβ accumulation in 3D hydrogel-based cultures due to limited diffusion of the Aβ peptides (*34, 38, 45*). High Aβ plaque load has been shown to induce pro-inflammatory responses in microglia from transgenic AD mouse models (*46, 47*). Hence, we next investigated whether MDMi are functional in 3D co-culture and respond to FITC-Aβ aggregates. We used a multiplex immunoassay to measure cytokines released in the conditioned medium of FITC-Aβ-treated 3D co-cultures, and 3D mono-cultures of ReNcell VM and MDMi. We observed a significantly upregulated secretion of classical pro-inflammatory cytokines IL-1β (2.8 fold) and IL-18 (2.3 fold), and similar trends for IL-6, in 3D co-cultures treated with FITC-Aβ compared to untreated conditions (**Fig. 2J**). Secretion of other pro-inflammatory cytokines, including IL-8, IFN-α2, MCP-1, IFN-γ and IL-10 was also stimulated by FITC-Aβ (**Fig. S4**). These inflammatory responses to Aβ were largely mediated by MDMi in the 3D co-cultures, as observed by significant changes in cytokine secretion following treatment in 3D MDMi mono-cultures, while 3D ReNcell VM mono-cultures remained unchanged (**Fig. 2J, S4**).

Overall, these results indicate that MDMi elicit a broad inflammatory response in the 3D platforms and demonstrate the functional capability of the 3D MDMi culture systems to model neuroinflammation in AD.

### Disease-specific differences in 2D and 3D mono-cultures of AD patient-derived MDMi

We next generated 2D and 3D MDMi mono-cultures from healthy control (HC) individuals and AD patients selected based on matched sex, age and APOE genotype (**Table 1**). Monocytes from HC and AD individuals were successfully differentiated into MDMi using 2D and 3D platforms (**Fig. 3A**). Survival of both HC and AD MDMi was significantly increased in 3D compared to 2D by 2.6 and 2.4 fold, respectively, while no differences in survival between HC and AD MDMi were observed in neither of the platforms (**Fig. 3B**).

**Fig. 3.**
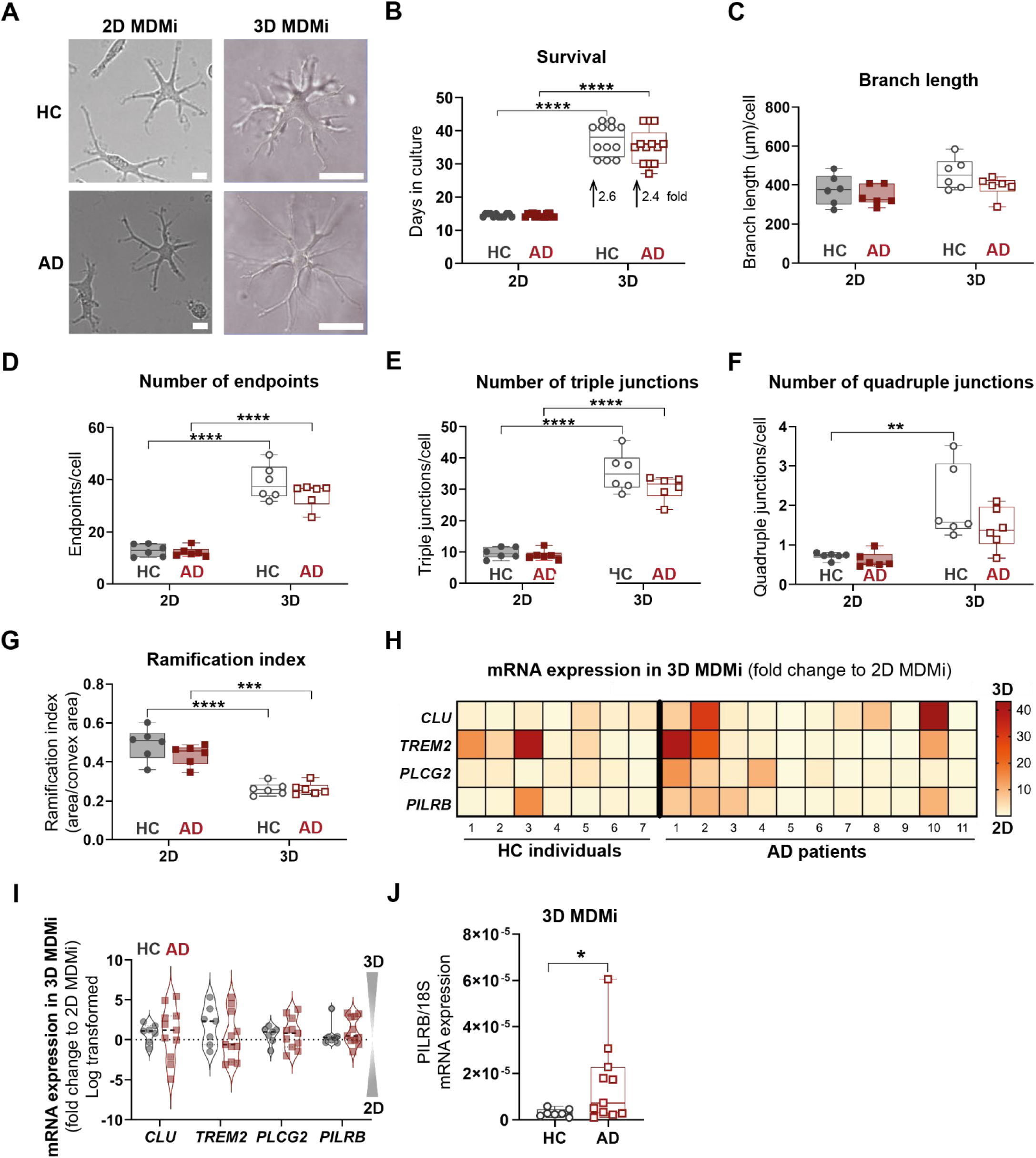
AD-associated phenotypes in 2D and 3D MDMi mono-cultures related to morphology and expression of AD risk genes. **(A)** Representative bright field images of HC and AD MDMi in 2D and 3D mono-cultures. Scale bars, 100 μm. **(B)** Survival of HC (*n* = 12) and AD (*n* = 13) MDMi in 2D and 3D mono-cultures. Quantification of morphological parameters in 2D and 3D MDMi from both HC (*n* = 6) and AD (*n* = 6) cohorts, including **(C)** branch length, **(D)** number of endpoints, **(E)** number of triple junctions, **(F)** number of quadruple junctions and **(G)** ramification index. **(H)** Heatmap representing HC (*n* = 7) and AD (*n* = 11) individual-specific fold changes in gene expression levels of the AD risk genes *CLU*, *TREM2*, *PLCG2* and *PILRB* in 3D MDMi compared to 2D. Red-yellow colour spectrum shows relative fold change of 3D MDMi as compared to 2D MDMi. **(I)** Violin plot representation of fold change (log transformed) of mRNA expression in 3D to 2D HC (*n* = 7) and AD (*n* = 11) MDMi. **(J)** mRNA expression of the AD risk gene *PILRB* in 3D HC (*n* = 7) and AD (*n* = 11) MDMi. Data are presented as mean ± SD. Each single data point represents one biological replicate. Two-way ANOVA with Tukey’s multiple comparison test; unpaired Student’s *t* test with or without Welch’s correction, two-tailed in **I**; \**P* < 0.05, \*\**P* < 0.01, \*\*\**P* < 0.001, \*\*\*\**P* < 0.0001.

To determine if 2D and 3D AD MDMi mono-cultures recapitulate disease-specific differences, we examined phenotypic features (morphology and expression of AD risk genes) associated with AD brain microglia in human and mouse models and compared them 1) between HC and AD cohorts and 2) between culture platforms. Quantification of morphological parameters (**Fig. S5**) revealed that when MDMi are cultured in 2D or 3D, HC and AD MDMi have similar branch length, number of branches (endpoints), number of triple and quadruple junctions, and ramification index (**Fig. 3C-G**). This suggests that HC and AD MDMi exhibit a similar branch structure and complexity in both 2D and 3D platforms.

We then assessed if those morphology parameters vary between culture platforms within the HC and AD cohorts. Comparison of branched structure parameters (branch length and number) between 2D and 3D MDMi showed similar trends in HC and AD, including similar branch length and increased number of branches in 3D compared to 2D (**Fig. 3C, D**). Interestingly, this contrasted with branch length in a young HC cohort (20-40 years of age) of MDMi (**Fig. 1E**), where MDMi exhibited longer branches in 3D compared to 2D. This may be explained by an age-related effect, where MDMi from elderly donors (60-80 years of age) have impaired response to branching signals and long-term culture in a 3D Matrigel scaffold. Further studies should confirm this observation.

Comparison of branch complexity parameters (triple and quadruple junctions) between 2D and 3D MDMi showed a similar trend in number of triple junctions but differed in number of quadruple junctions between the HC and AD cohorts. While the number of triple junctions increased in 3D compared to 2D MDMi in both HC and AD (**Fig. 3E**), the number of quadruple junctions increased in 3D compared to 2D MDMi in HC but remained unchanged in AD (**Fig. 3F**). Comparison of ramification index showed a significant decrease in 3D compared to 2D MDMi in both HC and AD cohorts, confirming a more ramified morphology in 3D MDMi (**Fig. 3G**). Overall, our results show that the 3D platform is able to enhance branch number, ramified morphology and complexity parameters in MDMi from HC individuals compared to 2D. However, the 3D platform induced an increase in branch number and ramified morphology but was not able to enhance all complexity parameters in MDMi from AD patients compared to 2D.

A recent study done in mouse models has suggested that AD risk genes present in microglia functionally influence microglial behaviour (*15*). In keeping with this, we examined 1) if risk genes are present/enriched in AD compared to HC MDMi, and 2) if these risk genes are differentially expressed in AD MDMi when cultured in 3D compared to 2D mono-cultures. A panel of AD risk genes including *CLU*, *TREM2*, *PLCG2* and *PILRB* was examined by qRT-PCR. Fold change of expression in 3D compared to 2D revealed heterogeneous distributions within the HC and AD cohorts, with 3D MDMi showing increased trends of enhanced expression within AD patients than HC individuals for most risk genes (*CLU*, *PLCG2* and *PILRB*) (**Fig. 3H, I**). Interestingly, *PILRB* expression was significantly upregulated in 3D AD MDMi compared to 3D HC MDMi (**Fig. 3J**), while no significant differences were observed for the other genes (**Fig. S6**). These results demonstrate that the 3D platform is able to enhance the expression of microglia-specific AD risk genes in AD MDMi compared to 2D and reflects the heterogeneity of disease phenotypes within AD patients.

### Disease-specific differences in 3D co-cultures of AD patient-derived MDMi

We next characterised MDMi from HC individuals and AD patients in the 3D co-culture platform (**Fig. 4A**). As observed in 3D MDMi mono-cultures, survival of MDMi in 3D co-culture was extended by 2.5 fold compared to 2D MDMi mono-cultures and was similar to 3D MDMi mono-cultures in both HC and AD cohorts (**Fig. 4B**). The increased survival of MDMi in 3D co-culture indicates that ReNcell VM do not affect MDMi viability. Consistently, similar expression of the apoptosis marker *BAX* between HC and AD 3D co-cultures (**Fig. 4C**) suggests that the cell ratio of MDMi and ReNcell VM and the duration of the co-culture were favourable.

**Fig. 4.**
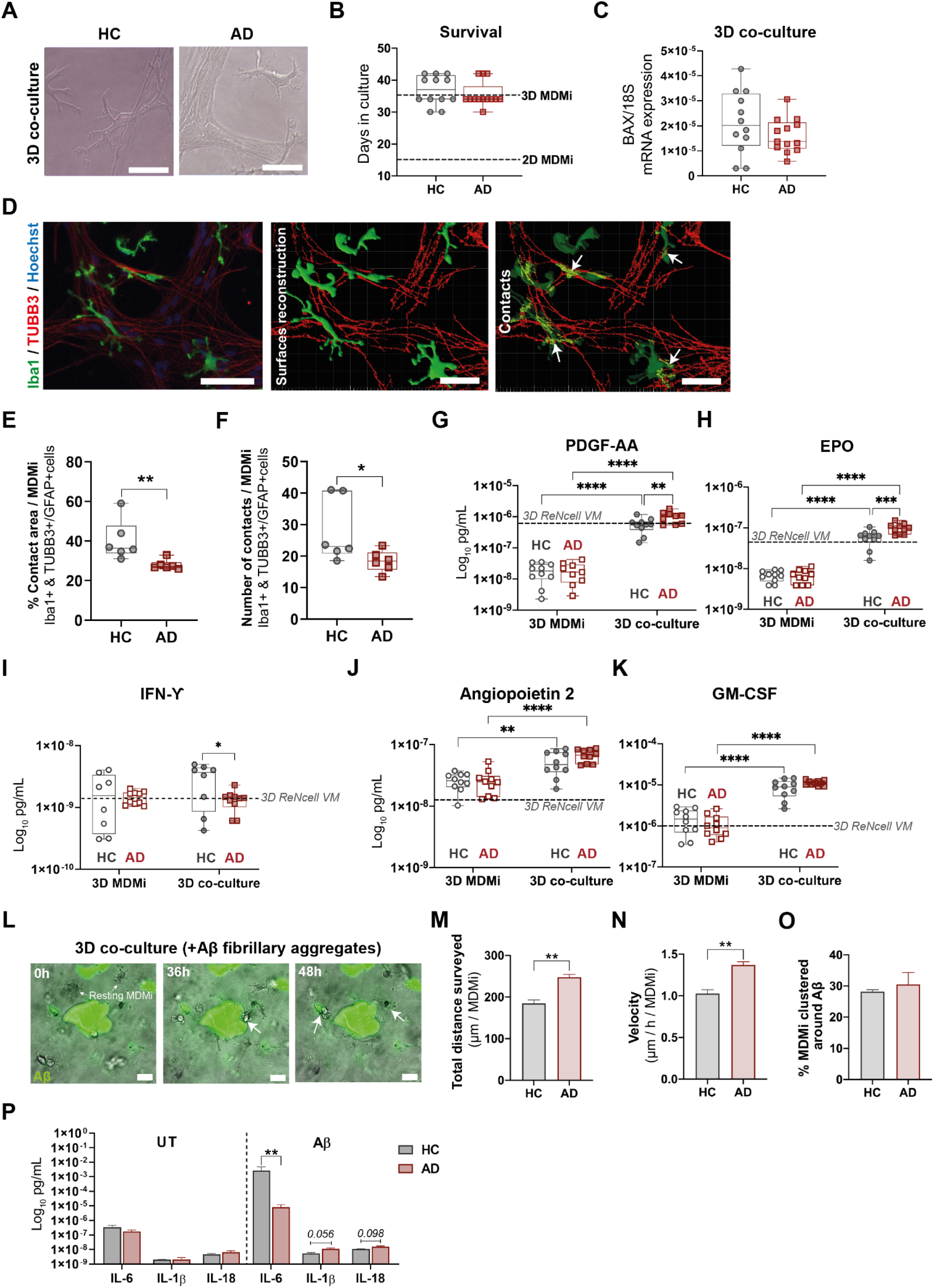
AD-associated phenotypes in 3D co-cultures related to cell-to-cell interaction with ReNcell VM, secretory activity and responses to Aβ aggregates. **(A)** Representative bright field images of HC and AD co-cultures. Scale bars, 100 μm. **(B)** Survival of HC (*n* = 12) and AD (*n* = 13) MDMi in 3D co-cultures. **(C)** Gene expression of the pro-apoptotic marker *BAX* in HC (*n* = 12) and AD (*n* = 13) 3D co-cultures. **(D)** Immunostaining against Iba1 (green) and TUBB3 (red) and 3D surface reconstruction of a 3D co-culture. Areas of contact between Iba1+ (MDMi) and TUBB3+ (ReNcell VM) cells are highlighted in yellow (white arrows). Scale bars, 100 μm. Quantification of **(E)** contact area and **(F)** number of contacts between Iba1+ and TUBB3+/GFAP+ cells in HC (*n* = 6) and AD (*n* = 6) 3D co-cultures. Concentration of **(G)** platelet-derived growth factors AA (PDGF-AA), **(H)** erythropoietin (EPO), **(I)** interferon-γ (IFN-γ), **(J)** angiopoietin 2 and **(K)** granulocyte-macrophage colony stimulating factor (GM-CSF) secreted by HC (*n* = 8-10) and AD (*n* = 9-10) MDMi in 3D mono-cultures and co-cultures. Baseline secretion by 3D ReNcell VM mono-cultures is represented with a dotted black line. **(L)** Representative images of 3D co-cultures containing FITC-Aβ aggregates in which MDMi exhibit a resting, ramified morphology at 0h and progressively become polarised acquiring an activated, round morphology with enlarged soma upon reaching and clustering on the Aβ deposit at 36 h and 48 h (white arrows). Scale bars, 100 μm. **(M)** Total surveillance distance, **(N)** velocity and **(O)** clustering around Aβ aggregates of HC (*n* = 2) and AD (*n* = 4) MDMi in 3D co-cultures containing Aβ aggregates. **(P)** Concentration of pro-inflammatory cytokines IL-6, IL-1β and IL-18 secreted by untreated (UT) and Aβ-treated HC (*n* = 3) and AD (*n* = 4) MDMi in 3D co-cultures. Data are presented as mean ± SD in **B**, **C**, **E-K**; mean ± SEM in **M-P**. Each single data point represents one biological replicate. Unpaired Student’s *t* test with or without Welch’s correction, two-tailed in **B**, **C**, **M-P**; Mann-Whitney test, two-tailed in **E**, **F**; two-way ANOVA with Tukey’s multiple comparison test in **G-K**; \**P* < 0.05, \*\**P* < 0.01, \*\*\**P* < 0.001, \*\*\*\**P* < 0.0001.

To investigate whether AD MDMi reflect disease-specific differences compared to HC MDMi in 3D co-culture, we analysed microglial behaviours known to be altered in AD brains, including 1) cell-to-cell interaction with neuro-glial cells, 2) secretion of growth factors and cytokines, and 3) migratory and inflammatory responses to Aβ aggregates. The marked synapse loss in AD is predominantly mediated by microglia through aberrant synapse engulfment (*6*). Hence, we first examined if the cell-to-cell interactions between AD MDMi and ReNcell VM show differences compared to HC MDMi in the 3D co-cultures. We performed a 3D rendering and subsequent surface reconstruction of immunofluorescent 3D co-culture images using the Imaris software (**Fig. 4D**). The *‘surface-surface contact area’* extension was used to quantify the area of contact between MDMi (labelled with Iba1) and ReNcell VM (labelled with the astrocyte- and neuron-specific markers GFAP and βIII-tubulin (TUBB3), respectively) and the number of contact points established between both cell types. Interestingly, we observed a significantly smaller area of contact and a reduced number of contact points between MDMi and ReNcell VM in AD compared to HC 3D co-cultures (**Fig. 4E, F**). This suggests an impairment in the cell-to-cell interactions between MDMi and ReNcell VM in AD 3D co-cultures that might have implications in disease.

In the context of AD, microglia secrete factors that alter neuron and astrocyte homeostasis, thereby contributing to disease pathogenesis (*48*). Hence, we next compared the secretory profiles of HC and AD MDMi in 3D co-cultures using a multiplex immunoassay in conditioned medium collected after 35 days of co-culture. Overall, we observed that AD MDMi in 3D co-culture secreted higher levels of platelet-derived growth factor AA (PDGF-AA) and erythropoietin (EPO) and lower levels of interferon-γ (IFN-γ) compared to HC MDMi in 3D co-culture (**Fig. 4G-I**). This indicates that AD MDMi exhibit an altered secretory activity in the 3D co-culture. Additionally, when comparing the secretion to 3D MDMi mono-cultures, we observed a significant upregulation of PDGF-AA, EPO (**Fig. 4G, H**) and other neurotrophic factors such as Angiopoietin 2 and the granulocyte-macrophage colony stimulating factor (GM-CSF) (**Fig. 4J, K**) in 3D co-cultures from both HC and AD cohorts. Secretion of these factors by 3D ReNcell VM mono-cultures was higher for PDGF-AA and EPO and lower for IFN-γ, Angiopoietin 2 and GM-CSF when compared to 3D MDMi mono-cultures (dotted lines in **Fig. 4G-K**). This suggests that the interaction between MDMi and ReNcell VM in the 3D co-cultures has a functional impact on either cell type. Overall, these results suggest that the 3D co-culture platform provides a suitable environment for AD MDMi to display functional impairments.

Microglia in the vicinity of Aβ plaques have been shown to exhibit altered proliferation, migration, clustering around Aβ aggregates and Aβ uptake in a mouse model of AD (*49*). Hence, in order to study MDMi behaviours in the presence of Aβ depositions, we added FITC-Aβ into 3D co-cultures from HC individuals and AD patients and live imaged for 7 days (**Fig. 4L**; **Movie S3**). We observed that AD MDMi surveyed longer distances and at a higher velocity around Aβ aggregates compared to HC MDMi, with no significant changes in the proportion of MDMi that clustered around the Aβ aggregates (**Fig. 4M-O**). Interestingly, measurement of inflammatory cytokine secretion using multiplex immunoassay revealed disease-specific differences between HC and AD MDMi in 3D co-cultures treated with FITC-Aβ. When Aβ was present in the cultures, IL-6 secretion was significantly decreased in AD compared to HC 3D co-cultures, while increasing trends were observed for IL-1β, IL-18 (**Fig. 4P**) and other pro-inflammatory cytokines such as TNF- α and IL-10 (**Fig. S7**). Such differences were not observed under untreated conditions. Together, these results indicate that AD MDMi respond differently to AD-related stressors compared to HC MDMi when modelled in the 3D co-culture platform.

### Drug treatment induces differential cytokine gene expression in MDMi cultured in 2D and 3D

Cytokines are key secreted molecules used by microglia to execute inflammatory and neuromodulatory functions. Moreover, altered cytokine levels have been reported in AD brains and may have important roles in disease pathogenesis (*50*). To investigate cytokine expression profiles in MDMi we analysed a panel of inflammatory cytokines (*IL-6*, *TNF-α*, *IL-8*, *TGF-β*, *IL-10, IL-1β* and *IL-18*) by qRT-PCR. We then examined for potential differences 1) between culture platforms and 2) between HC and AD cohorts. When comparing cytokine expression levels between platforms in the HC cohort, no significant differences were observed for any of the cytokines (**Fig. 5A**). However, in the AD cohort, *IL-8* was significantly downregulated in 3D co-culture compared to 3D MDMi mono-culture, and *TGF-β* and *IL-18* were significantly downregulated in 3D co-culture compared to 2D MDMi (**Fig. 5B**). When comparing between HC and AD MDMi in either platform, we observed a significantly decreased expression of *TNF-α* in AD MDMi compared to HC only in the 3D MDMi platform (**Fig. S8A**). No disease-specific differences were observed for the rest of cytokines (**Fig. S8B-G**). Together, these results demonstrate cytokine changes dependent on platform, which were more evident in MDMi from AD patients than HC individuals.

**Fig. 5.**
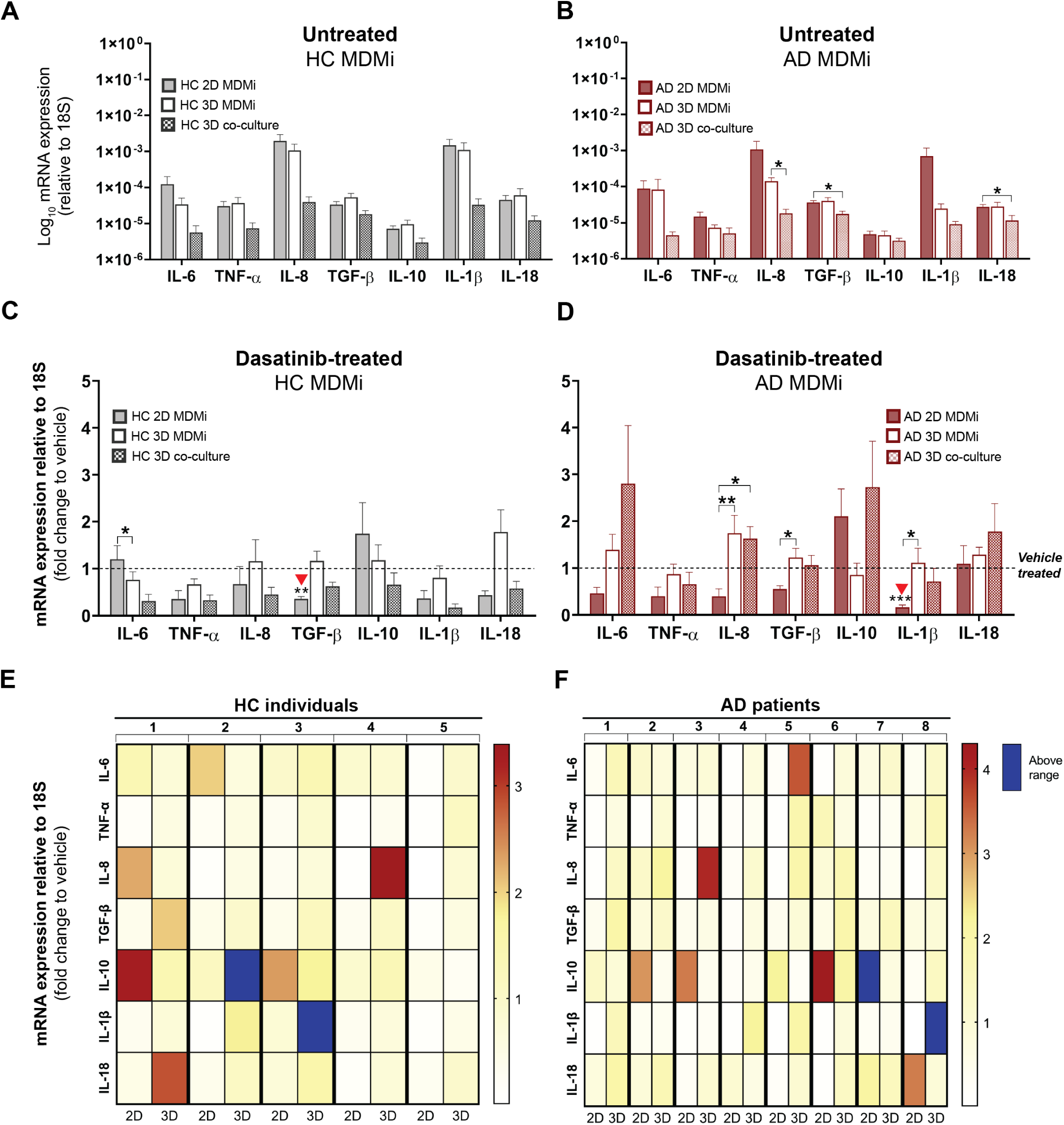
Dasatinib treatment induces cytokine expression responses that differ between culture format and are heterogeneous among HC and AD MDMi. Log-transformed mRNA expression of the inflammatory cytokines IL-6, TNF-α, IL-8, TGF-β, IL-10, IL-1β and IL-18 in untreated **(A)** HC (*n* = 8) MDMi and **(B)** AD (*n* = 12) MDMi in all culture formats (2D and 3D mono-cultures and 3D co-cultures). Fold change in cytokine mRNA expression levels following 24 h exposure to 100nM dasatinib compared to vehicle (DMSO)-treated cultures in **(C)** HC (*n* = 6) and **(D)** AD (*n* = 8) 2D and 3D MDMi mono-cultures and 3D co-cultures. Red arrowheads indicate significance compared to vehicle-treated condition. Dotted black lines represent baseline responses of vehicle-treated cultures. Heatmaps showing **(E)** HC (*n* = 5) and **(F)** AD (*n* = 8) donors-specific changes in mRNA expression from MDMi mono-cultures in 2D and 3D. Red-yellow colour spectrum represents relative fold change of mRNA expression after dasatinib treatment compared to vehicle. Expression changes falling outside the displayed range are indicated in dark blue. Data are presented as mean ± SEM. One-way ANOVA with Dunnett’s multiple comparison test; \*\**P* < 0.01, \*\**P* < 0.01, \*\*\**P* < 0.001.

Based on the differences in baseline cytokine expression between platforms, we next investigated whether drug treatment alters such platform-dependent responses in the HC and AD cohorts. For this, we trialled two FDA-approved compounds, dasatinib and spiperone, which have anti-cancer and anti-psychotic properties, respectively. Additionally, these drugs have been shown to mitigate inflammation in *in vitro* models of microglia (*51, 52*) and murine models of AD (*53*) and have potential as re-purposed drugs for treating neuroinflammation. Dasatinib-treated MDMi from HC individuals showed a significantly reduced expression of *IL-6* in 3D compared to 2D MDMi, while expression levels for the rest of cytokines remained unchanged between platforms (**Fig. 5C**). Dasatinib-treated MDMi from AD patients showed a significantly increased expression of *IL-8*, *TGF-β* and *IL-1β* in 3D compared to 2D MDMi. Similarly, *IL-8* was upregulated in 3D co-cultures compared to 2D MDMi (**Fig. 5D**). Spiperone-treated MDMi showed similar cytokine expression levels in all platforms in the HC cohort (**Fig. S9A**) and a significant downregulation of *TNF-α* in 3D AD co-cultures compared to 3D AD MDMi mono-cultures (**Fig. S9B**). When compared to untreated conditions, dasatinib induced significant changes (*i.e.,* downregulation of *TGF-β* and *IL-1β* in HC and AD MDMi, respectively) only in the 2D platform (**Fig. 5C, D**), while spiperone did not significantly alter cytokine expression in any platform (**Fig. S9A, B**).

Interindividual variability in drug responses was displayed using heatmaps, which show cytokine expression levels in 2D and 3D MDMi from each individual in the HC and AD cohorts (**Fig. 5E, F**; **Fig. S9C, D**). When comparing within HC individuals or AD patients, cytokine expression in MDMi showed high heterogeneity in both 2D and 3D platforms. For example, in the dasatinib-treated HC MDMi cohort, *IL-10* was highly expressed in 2D MDMi from individuals 1 and 3 compared to individuals 4 and 5 (**Fig. 5E**). In the dasatinib-treated AD MDMi cohort, *IL-10* was highly expressed in 2D MDMi from patients 2, 3, 6 and 7 compared to patients 4 and 8 (**Fig. 5F**). When considering dasatinib-treated 3D MDMi, *IL-8* showed highly variable expression in HC and AD, with individual 4 and patient 3 showing a notably upregulated gene expression compared to the rest of patients (**Fig. 5E, F**). Importantly, differences between dasatinib-treated 2D and 3D MDMi in each particular HC or AD individual were also demonstrated in the heatmaps. Some examples include divergent expression levels of *IL-8* in HC individual 4, *IL-10* in HC individual 1 (**Fig. 5E**), *IL-8* in AD patient 3 and *IL-10* in AD patients 2 and 3 (**Fig. 5F**). Similar trends in the heterogeneity of drug responses within and between 2D and 3D MDMi were observed for spiperone-treated MDMi (**Fig. S9C, D**).

Overall, these results suggest that drug treatment induces differential cytokine expression in MDMi from both HC individuals and AD patients in a culture platform-dependent manner.

## Discussion

Research involving human microglia is hampered by the lack of model systems that faithfully recapitulate their dynamic characteristics *in vivo*, particularly those associated with disease. Generating more representative microglia model systems will address this shortcoming and help elucidate microglia-mediated disease mechanisms, expediting pre-clinical investigations of microglia-targeted therapeutics and better clinical outcomes (*54*).

Recreating physiologically relevant culture conditions to mimic the interaction of microglia with other brain cell types and the extracellular matrix is crucial for accurately modelling the role of microglia in disease. To date, no study has attempted to culture human microglia that have not been genetically modified in a 3D system resembling human brain tissue. In an effort to develop more representative *in vitro* models of human microglia, we used MDMi, a model system of microglia-like cells that has emerged as a promising, patient-specific drug screening platform for neurological diseases (*55, 56*). Previous reports have shown that MDMi morphologically and functionally resemble brain-resident human microglia and express bonafide microglial markers (*28–31*). In this study, we developed novel MDMi platforms that incorporate relevant *in vivo* cues resembling the microenvironment of the brain. This was achieved by utilising a hydrogel-based 3D model to culture MDMi in microenvironments of increasing complexity and physiological relevance, firstly as 3D MDMi mono-cultures and secondly as 3D co-cultures of MDMi with neural progenitor cells.

Our study showed that a 3D hydrogel-based culture enhances MDMi survival, the extent and complexity of ramification (branching) and the expression of mature microglial markers (*i.e., TMEM119*) compared to standard 2D culture conditions. In addition, we have demonstrated the feasibility of culturing MDMi together with neuro-glial cells derived from human immortalised neural progenitor cells (*i.e.,* ReNcell VM) in a 3D co-culture setting. Our findings also confirmed that MDMi are capable of producing broad inflammatory responses upon exposure to inflammatory stimuli, which is preserved in both 3D mono- and co-culture models.

We observed an enhanced maturation of microglial and neuro-glial cells in both our MDMi and ReNcell VM 3D mono-cultures compared to 2D, in keeping with previous studies that used similar 3D cell models of primary rodent microglia and astrocytes (*57, 58*) and human induced neural stem cell lines (*59, 60*). The enhanced maturation of microglia-like features in 3D MDMi could be attributed to their increased survival in culture. Our model therefore shows two major advantages. Firstly, it better mimics *in vivo* growth conditions. Secondly, it provides a longer time frame for differentiation of monocytes into more mature MDMi that more closely represent brain microglia. Future studies should examine whether MDMi in 3D co-culture develop an enhanced maturation of microglia-like features compared to MDMi in 2D and 3D mono-cultures.

Microglia are involved in the pathogenesis of AD and contribute to the clinical heterogeneity observed among patients. Hence, 3D MDMi models offer a great opportunity to study AD microglia in a patient-specific manner. To investigate the capability of MDMi to model phenotypical features of AD microglia in the 2D and 3D mono-culture and co-culture platforms, we generated MDMi cultures using monocytes from living AD patients, a major advantage of our 3D AD MDMi platform. Unlike murine or human immortalised microglia cell lines used in previous studies (*45, 61*), MDMi are patient-derived and have not been genetically modified, being therefore more physiologically relevant and clinically applicable. In addition, the use of cell samples obtained from living patients allows for longitudinal modelling of disease progression in AD, a prerequisite for targeted treatment at various stages of the disease.

We first compared cellular aspects in our 2D and 3D MDMi mono-culture models that have been reported to be altered in microglia isolated from post-mortem AD patient brains. Analysis of the MDMi branched morphology revealed similarities between HC and AD MDMi in both the 2D and the 3D mono-culture models. These similarities most likely reflect age-related phenotypes described in human AD microglia rather than activated phenotypes observed in microglia from AD mouse models. Indeed, this age-related phenotype, termed HAM (human Alzheimer’s microglia) was originally described in myeloid cell populations isolated from human AD post-mortem brain samples at later stages of disease (*11*). The HAM profile showed substantial overlap of age-associated gene expression patterns between AD and control samples. Therefore, HC and AD MDMi are likely to have similar morphologies irrespective of the culture platform. Our 3D platform revealed a failure of AD MDMi to fully increase morphological complexity, particularly in relation to the number of quadruple junctions (points connecting four sub-branches), compared to 2D. However, all complexity parameters, including number of triple (points connecting three sub-branches) and quadruple junctions, were enhanced in 3D HC MDMi compared to 2D. This potentially suggests that AD MDMi have an altered branching complexity, only evident when culturing in a 3D platform. Microglial branches are highly dynamic and constantly survey the brain microenvironment, phagocytosing dysfunctional synapses and releasing trophic factors that support neural connectivity (*62*). Prior studies have identified a reduced number of junctions in human AD microglial branches (*63*) and less AD microglial arborisation area compared to age-matched HC (*64*). Our findings highlight that despite longer differentiation time in a brain-like 3D structural environment, AD MDMi in 3D are not able to fully increase the outgrowth of sub-branches in the same way as HC MDMi.

Gene expression of microglial AD risk genes, including *CLU*, *TREM2*, *PLCG2* and *PILRB*, in AD MDMi, revealed an overall increased expression of AD risk genes in 3D compared to 2D mono-cultures. However, there was prominent person-to-person variation in the expression level of the above genes within both HC and AD cohorts, which reflects clinical heterogeneity observed in patients. Importantly, *PILRB* (an activating immune receptor) was significantly upregulated in AD MDMi 3D mono-cultures compared to HC, which suggests that the 3D MDMi model provides a suitable screening platform with improved clinical translation to identify other AD risk genes and microglia-targeted therapeutics.

To assess the fidelity of AD MDMi in recapitulating disease-specific alterations in 3D co-cultures, we evaluated aspects such as the interaction of MDMi with neuro-glial cells, their secretory profiles and migration around Aβ aggregates. As previously described, microglia establish physical contacts with neurons through identified molecular mechanisms (*65*). Our results showed reduced physical contacts between AD MDMi and ReNcell VM-derived neuro-glial cells compared to HC. Alterations in such microglia-neuron interactions could impact the microglial capacity to respond to neuronal damage, providing a potential mechanism underlying neuron degeneration in AD.

Microglia in AD and ageing have dysfunctional secretomes (*66*). We observed altered secretion patterns of PDGF-AA, EPO and IFN-γ in AD 3D co-cultures when compared to HC, in agreement with previous observations in AD patients. An unbalanced distribution of PDGF-AA immunopositive cells has been linked to gliosis in AD post-mortem brains (*67*). Moreover, protein expression levels of EPO receptors in astrocytes have been found to be altered in post-mortem hippocampal brain sections from AD patients (*68*). Likewise, plasma levels of IFN-γ have been shown to fluctuate according to disease severity in AD patients (*69*) and studies in mice have demonstrated that IFN-γ impacts neurogenesis and synaptic plasticity (*70*). Taken together, our findings suggest that MDMi in 3D co-culture recapitulate alterations reported in AD patients and are therefore an exciting new platform for disease modelling.

Microglia have a canonical role in the removal of Aβ aggregates (*71*). In order to validate the functional response of MDMi to Aβ aggregates, we investigated the behaviour of these cells in 3D co-culture in the presence of Aβ aggregates. Our results showed that AD MDMi migrate longer distances at a faster speed compared to HC. Similarly, elevated migration rates were observed in human immortalised microglia cultured in a 3D tri-culture model with ReNcell VM overexpressing pathogenic Aβ species (*45*). However, an AD mouse model with aberrant Aβ production showed decreased microglial migration towards Aβ plaques (*72*), highlighting important differences between human and murine microglia responses to Aβ. Our findings warrant further investigation of microglial motility as a potentially dysregulated cellular feature in AD brains. Whether an increase in surveillance and speed of AD MDMi correlates with an impaired branched complexity in these cells remains to be elucidated in future studies.

Interestingly, we did not observe differences in the number of MDMi that clustered around Aβ aggregates. However, we observed changes in pro-inflammatory cytokine secretion in MDMi from AD compared to HC. Altogether, this suggests that AD MDMi may have unique disease-specific chemotactic and secretory responses against Aβ. Future studies should investigate how such changes in MDMi impact the phagocytic clearance of Aβ aggregates in the 3D co-cultures. Overall, disease-specific differences exhibited by AD MDMi in the 3D co-culture platform confirm the possibility to model disease in AD patient-specific MDMi using culture platforms that better recapitulate the brain microenvironment.

Preliminary drug testing demonstrated the utility of the 3D MDMi models as personalised drug screening tools. The main reasons are as follows. First, differences in MDMi drug responses between 2D and 3D culture conditions reflect the functional impact of MDMi cultured in a more biologically relevant 3D environment. This is in agreement with a previous study carried out on tumour cell lines, which described varied treatment outcomes relating to cell proliferation depending on the culture system (*73*). Moreover, another study reported more similarities in drug-induced cellular responses between 3D cultures and *in vivo* conditions than compared to 2D cultures (*74*). Future investigations should determine whether drug responses from our 3D MDMi models correlate better with responses identified in animal models and clinical data from patients. Second, patient heterogeneity in MDMi drug responses was evident, supporting the translatability of our 3D platforms to measure individual patient responses in the clinic and further validating our *in vitro* systems as promising alternative platforms for personalised drug screening.

To further enhance our 3D MDMi model systems, microfluidic technology that accounts for the dynamic interstitial fluid flow within the brain could be used (*75*). Moreover, the addition of patient hiPSC-derived neural progenitor cells into the 3D MDMi co-cultures would make this platform more personalised and likely a more accurate representation of the human brain. Finally, more-defined synthetic hydrogels may enable us to more carefully dissect the functionality of MDMi in 3D cultures given their consistent and tuneable properties.

In conclusion, we describe reproducible and easy-to-generate 3D *in vitro* models of MDMi that are able to recapitulate potentially important AD-specific differences associated with diseased microglia, not identified in 2D models. This study opens new doors to generate patient-specific drug testing platforms that support the development of microglia-targeted therapeutic interventions tailored for AD patients and potentially other neurological disorders.

## Acknowledgements

We would like to thank the volunteers from QIMR Berghofer who have donated blood to this project and the participants recruited through the PISA study. We also thank the Microscopy, Flow cytometry, Sample Processing and Statistical teams at QIMR Berghofer for their assistance. Special thanks to Kurt Giulani for helpful insights on LEGENDplex^TM^ data analysis. Lastly, we acknowledge the coordinators of the PISA study, Professor Michael Breakspear, Jessica Adsett and Natalie Garden.

## Author contributions

C.C-L., H.Q., L.E.O. and A.R.W. conceived and designed the study. C.C-L. and H.Q. performed experiments. C.C-L., H.Q. and T.H.N. analysed data. C.C-L., H.Q., R.S. and A.R.W. interpreted data. Y.S., C.C.G. and M.K.L. coordinated blood collection from the PISA study. C.C-L. wrote the manuscript. H.Q., R.S., L.E.O. and A.R.W. provided critical feedback in reviewing and editing. All authors reviewed and commented on the final version of the manuscript.

---

**Table S1.**
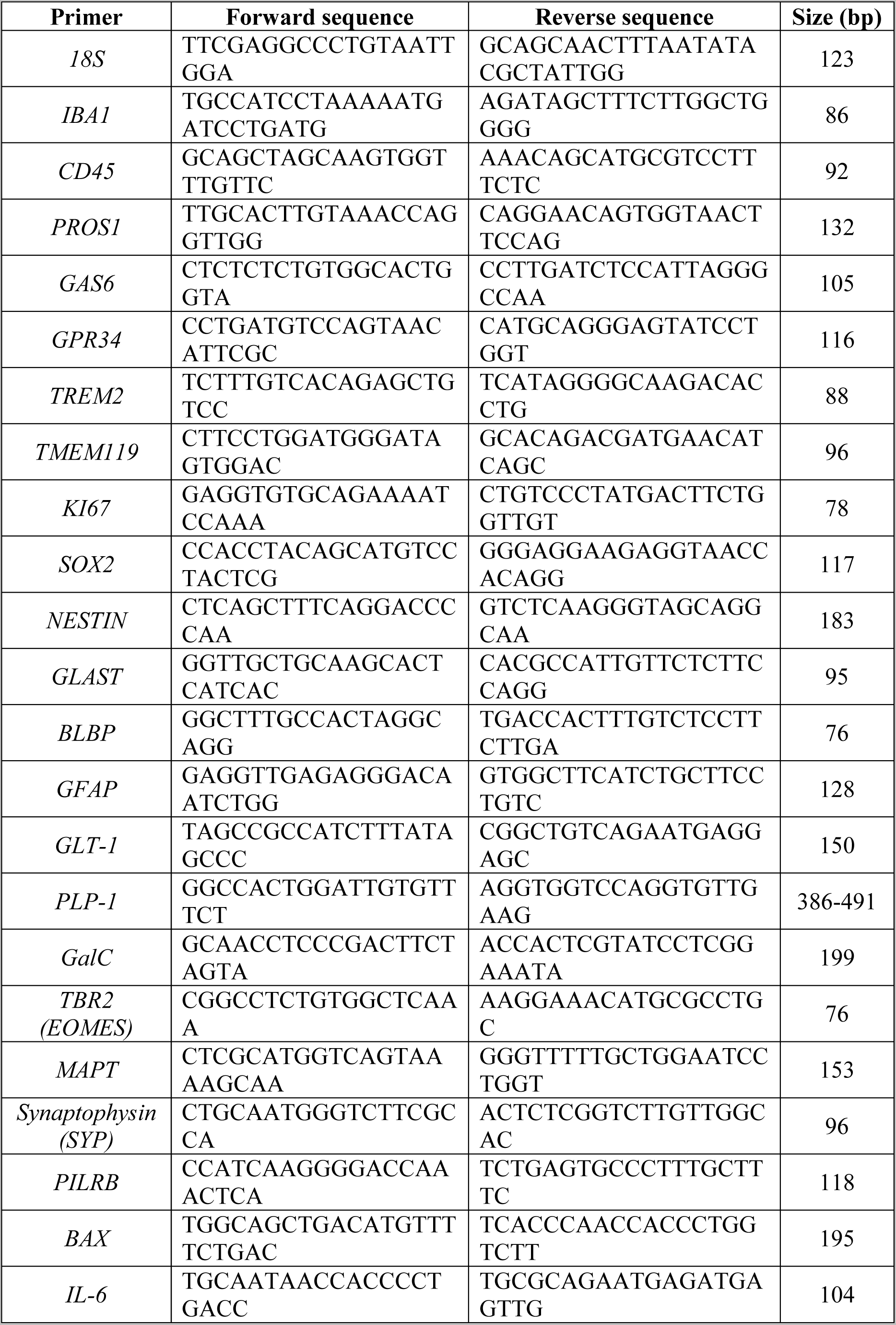
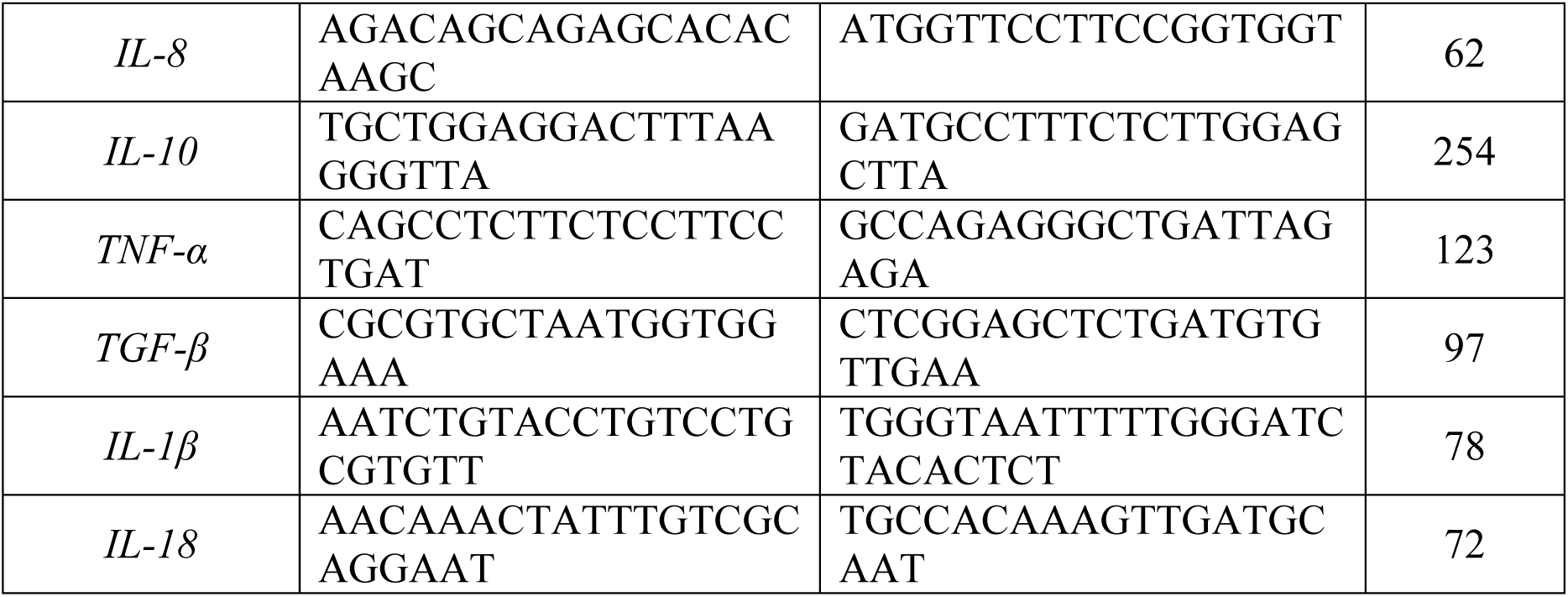
qRT-PCR primer sequences.

**Fig. S1.**
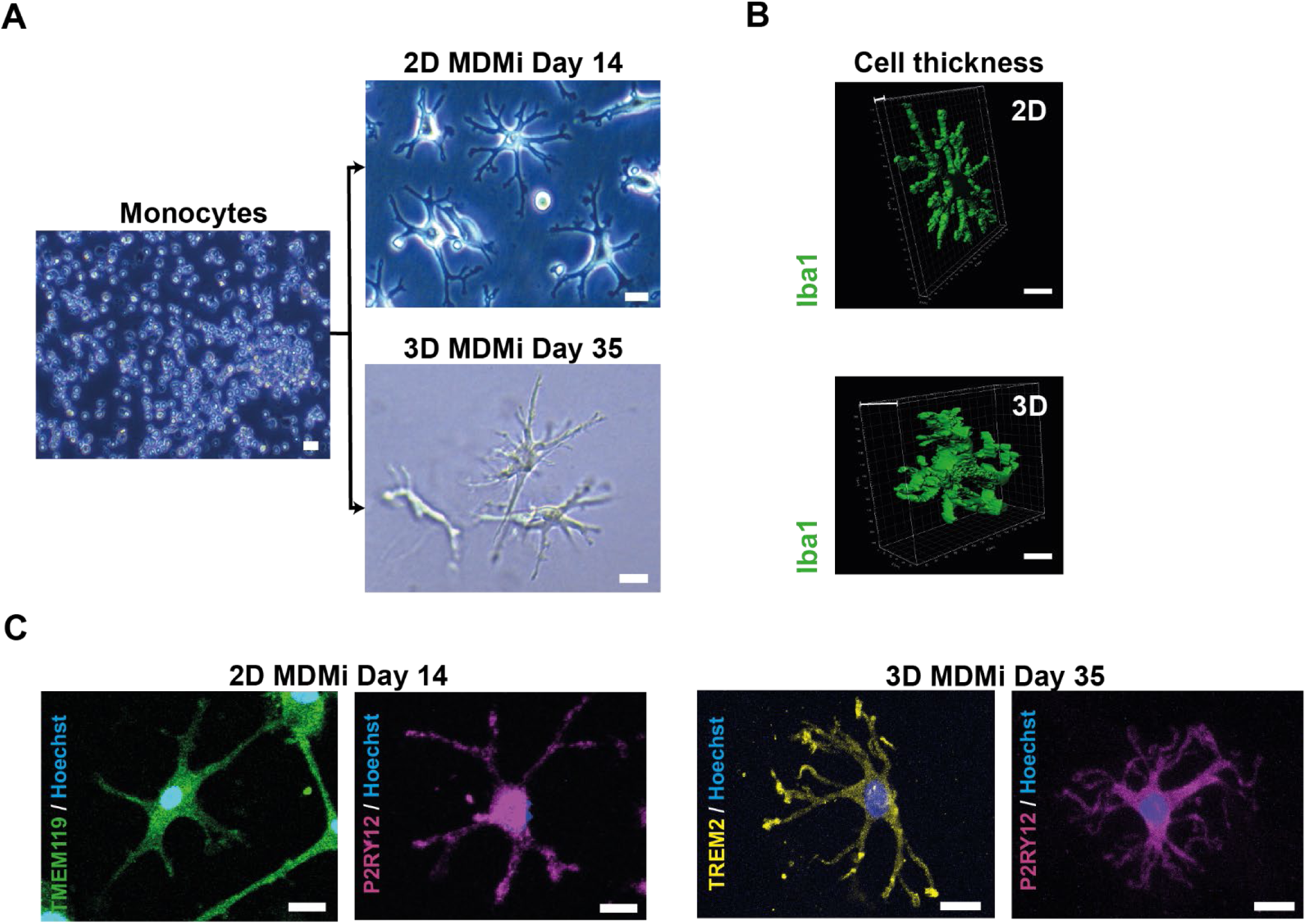
2D and 3D MDMi show different cell thickness and positive immunostaining for microglia-enriched markers. **(A)** Representative bright field images of monocytes and 2D and 3D MDMi after 14 (top) or 35 (bottom) days of differentiation, respectively. Scale bars, 25 μm. **(B)** 3D reconstruction images of Iba1-stained MDMi and thickness of 2D and 3D MDMi. Scale bars, 25 μm. **(C)** Immunofluorescence of 2D and 3D MDMi for TMEM119, P2RY12 and TREM2. Scale bars, 25 μm.

**Fig. S2.**
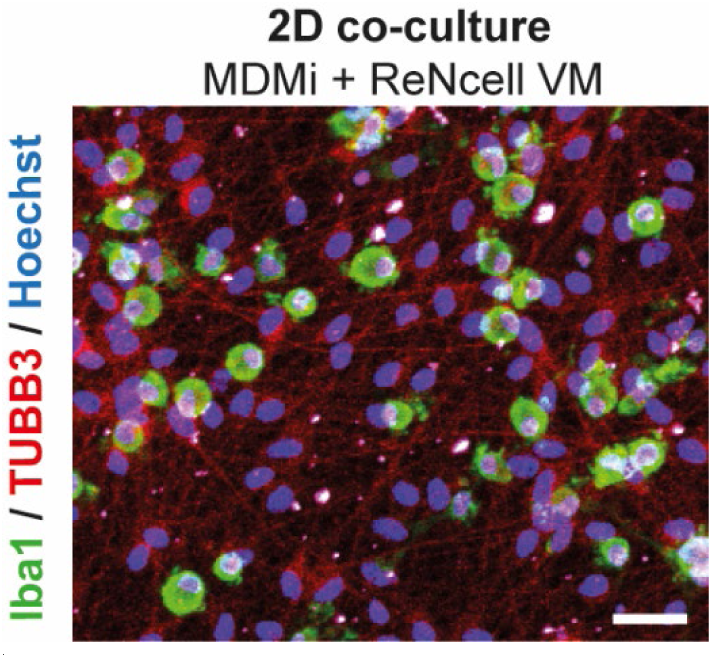
Characterisation of 2D MDMi and ReNcell VM co-cultures. MDMi co-culture with ReNcell VM-derived neuro-glial cells in 2D is insufficient for monocyte differentiation into MDMi, as monocytes retain a round morphology after 40 days in 2D co-culture. MDMi were stained for Iba1 and ReNcell VM were stained for β3-tubulin (TUBB3). Scale bar, 100 μm.

**Fig. S3.**
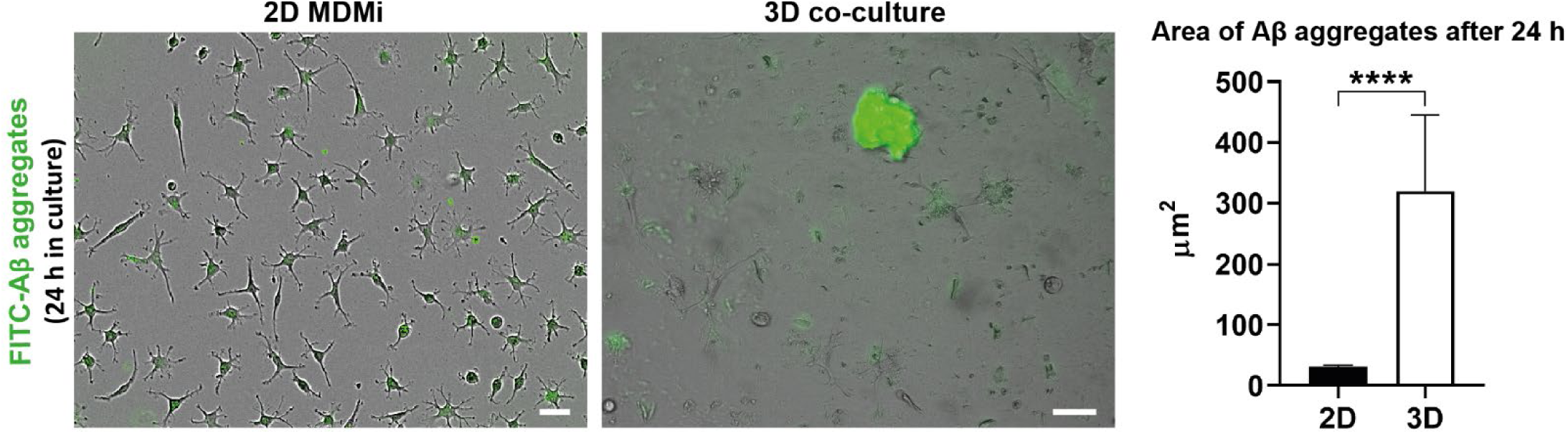
FITC-Aβ peptides form larger aggregates in 3D compared to 2D after 24 h in culture. FITC-Aβ peptides were added to 2D MDMi and 3D co-cultures. Imaging was conducted following incubation for 24 h. Area of Aβ aggregates (µm^2^) was quantified for comparison between the 2D MDMi (*n* = 3) and 3D co-cultures (*n* = 3). Scale bars, 100 μm. Data are presented as mean ± SEM. Mann-Whitney test, two-tailed; \*\*\*\**P* < 0.0001.

**Fig. S4.**
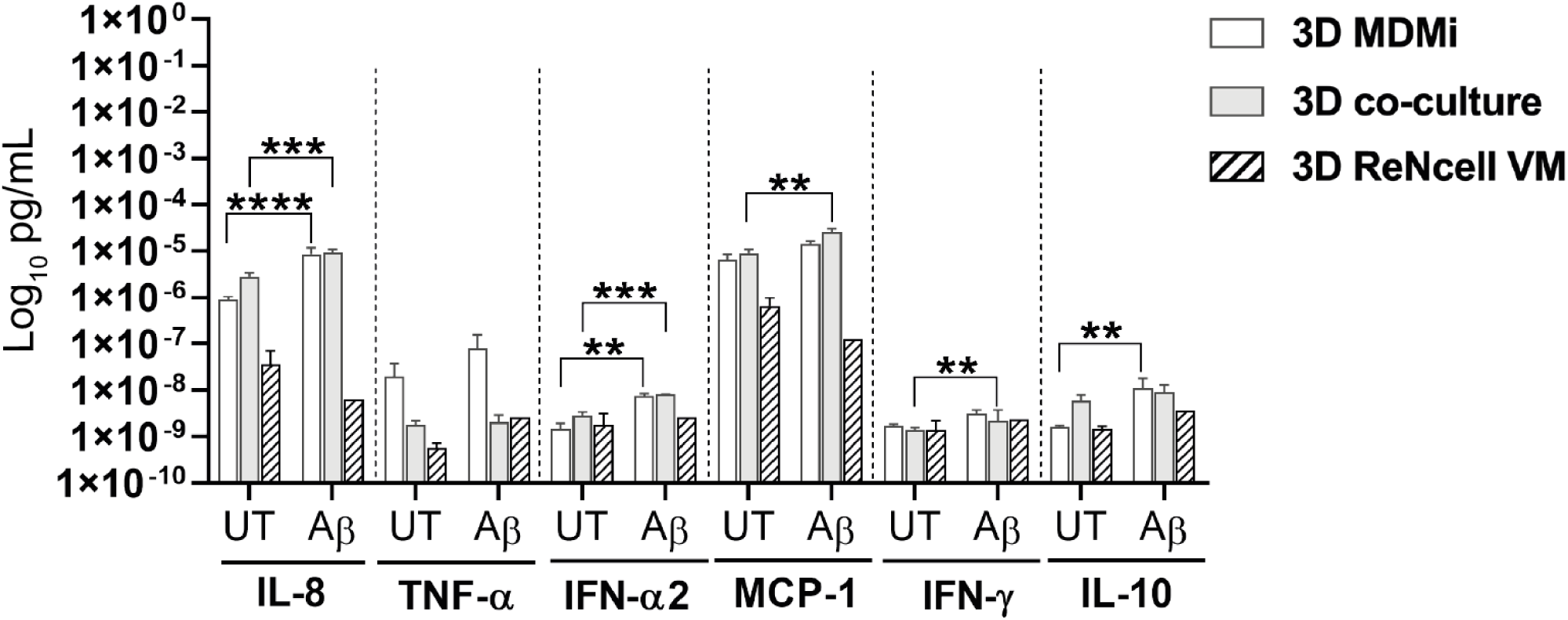
Pro-inflammatory cytokine secretion profiles of 3D MDMi, 3D co-culture and 3D ReNcell VM. Concentration of secreted pro-inflammatory cytokines IL-8, TNF-α, IFN-α2, MCP-1, IFN-γ and IL-10 by 3D MDMi (*n* = 2) and ReNcell VM (*n* = 1) mono-cultures and 3D co-cultures (*n* = 3) upon exposure to FITC-Aβ aggregates for 7 days. Data are presented as mean ± SEM. Unpaired Student’s *t* test with or without Welch’s correction, two-tailed; \*\**P* < 0.01, \*\*\**P* < 0.001, \*\*\*\**P* < 0.0001.

**Fig. S5.**
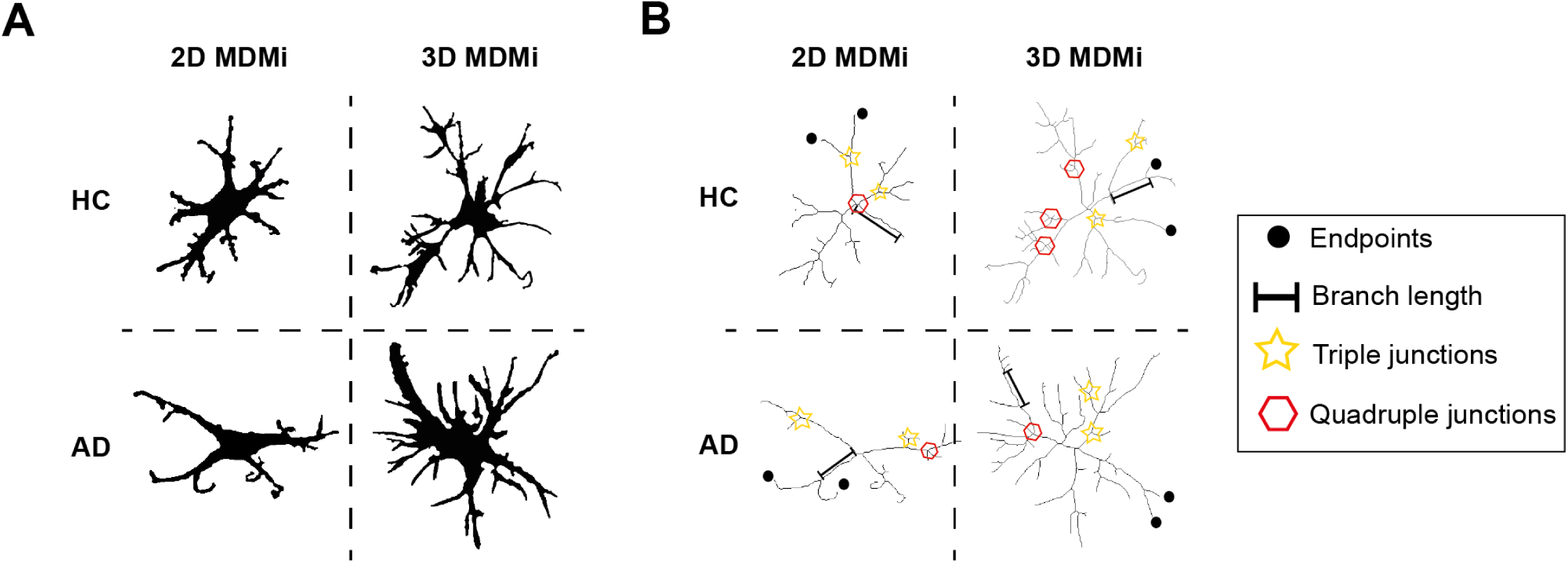
Depiction of morphological parameter measurements in HC and AD MDMi in 2D and 3D mono-cultures. **(A)** Binary and **(B) s**keleton images of HC and AD MDMi in 2D and 3D mono-cultures showing the morphological parameters (endpoints, branch length, triple and quadruple junctions) analysed to estimate the branched structure and complexity of MDMi.

**Fig. S6.**
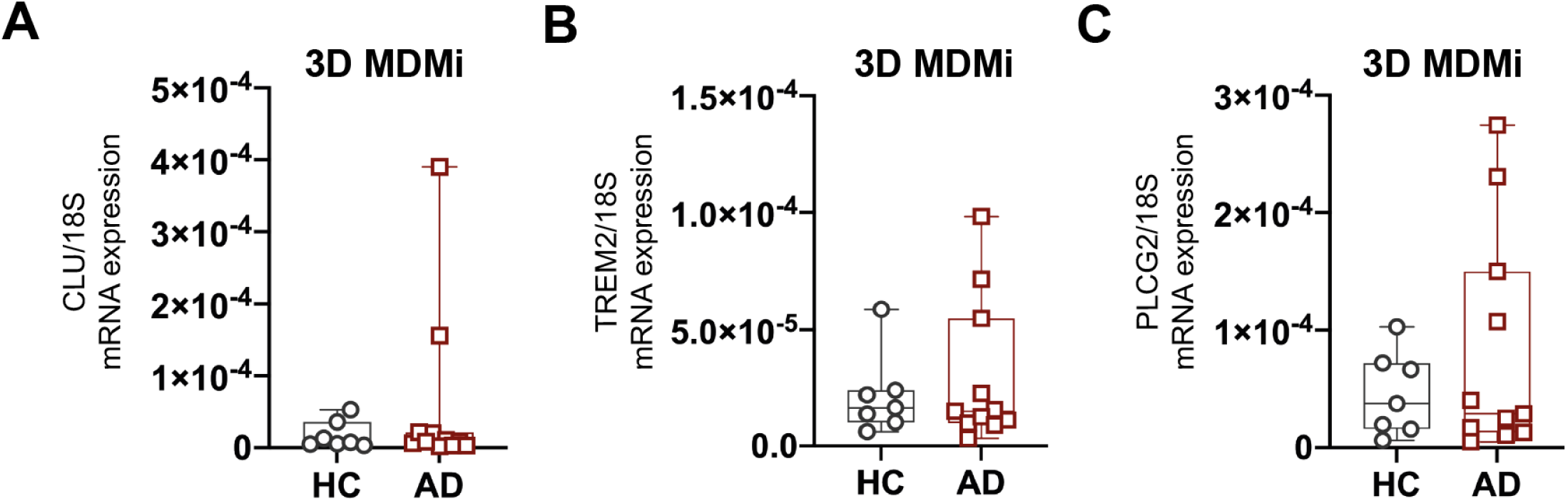
Gene expression of AD risk genes in HC and AD 3D MDMi. qRT-PCR quantification of **(A)** *CLU*, **(B)** *TREM2* and **(C)** *PLCG2* mRNA expression in 3D MDMi mono-cultures from HC (*n* = 6) and AD (*n* = 11) donors. Data are presented as mean ± SD. Each single data point represents one biological replicate. Unpaired Student’s *t* test with or without Welch’s correction, two-tailed.

**Fig. S7.**
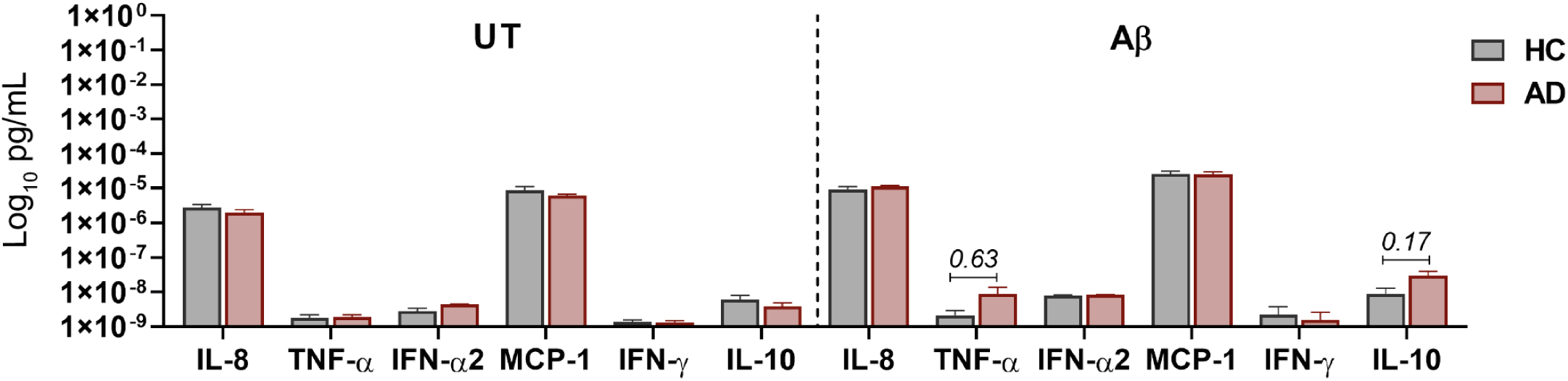
Inflammatory cytokine secretion by HC and AD 3D co-cultures under untreated and Aβ-treated conditions. Concentration of secreted IL-8, TNF-α, IFN-α2, MCP-1, IFN-γ and IL-10 are displayed in HC (*n* = 3) and AD (*n* = 4) MDMi 3D co-cultures. Data are presented as mean ± SEM. Unpaired Student’s *t* test with or without Welch’s correction, two-tailed.

**Fig. S8.**
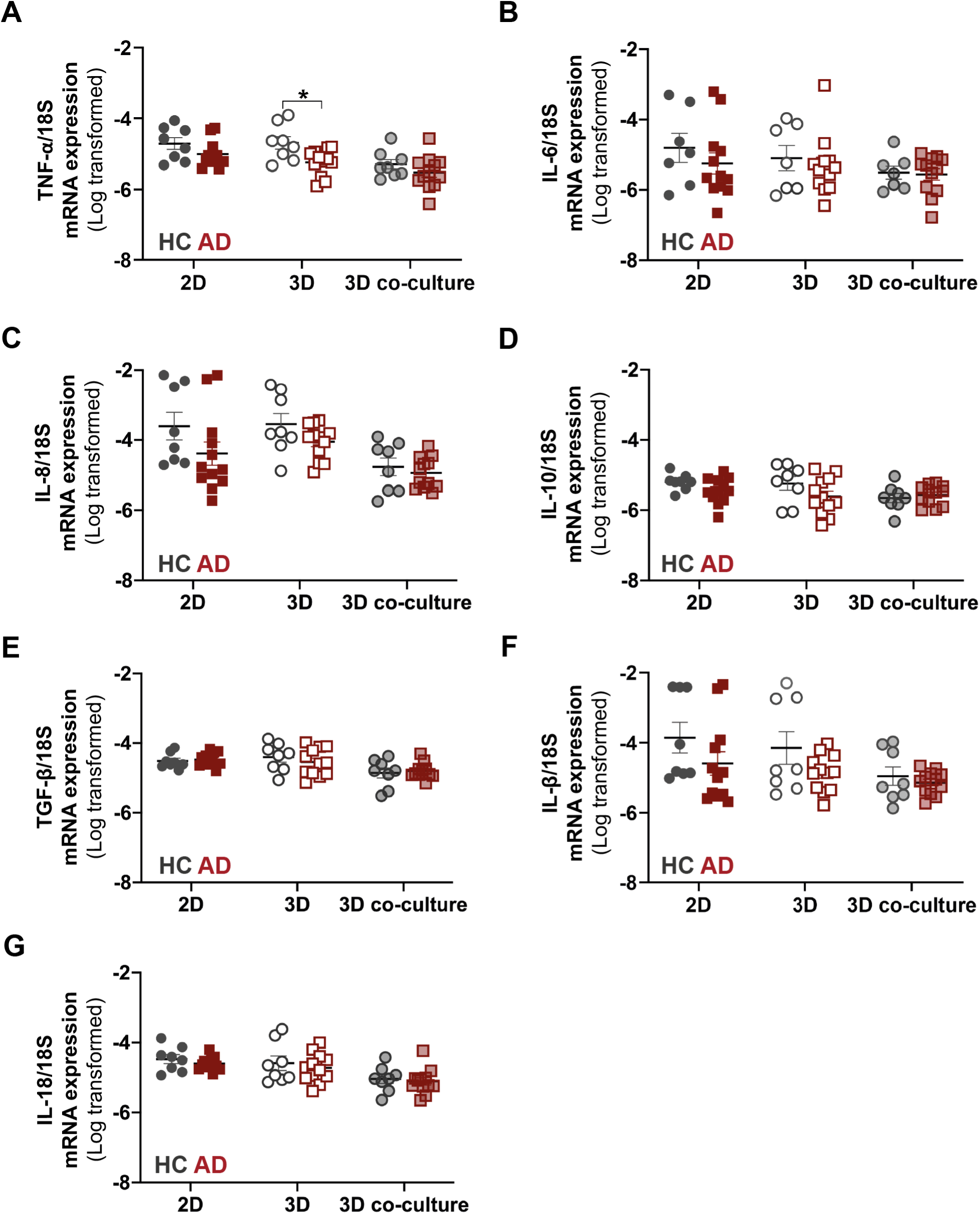
Cytokine expression profiles in HC and AD MDMi 2D and 3D mono-cultures and 3D co-cultures. Log-transformed mRNA expression of the inflammatory cytokines **(A)** TNF-α, **(B)** IL-6, **(C)** IL-8, **(D)** IL-10, **(E)** TGF-β, **(F)** IL-1β and **(G)** IL-18 in HC (*n* = 7-8) and AD (*n* = 12) MDMi cultures. Data are presented as mean ± SD. Each single data point represents one biological replicate. Comparisons between HC and AD MDMi in either culture format were performed using unpaired Student’s *t* test with or without Welch’s correction, two-tailed; \**P* < 0.05.

**Fig. S9.**
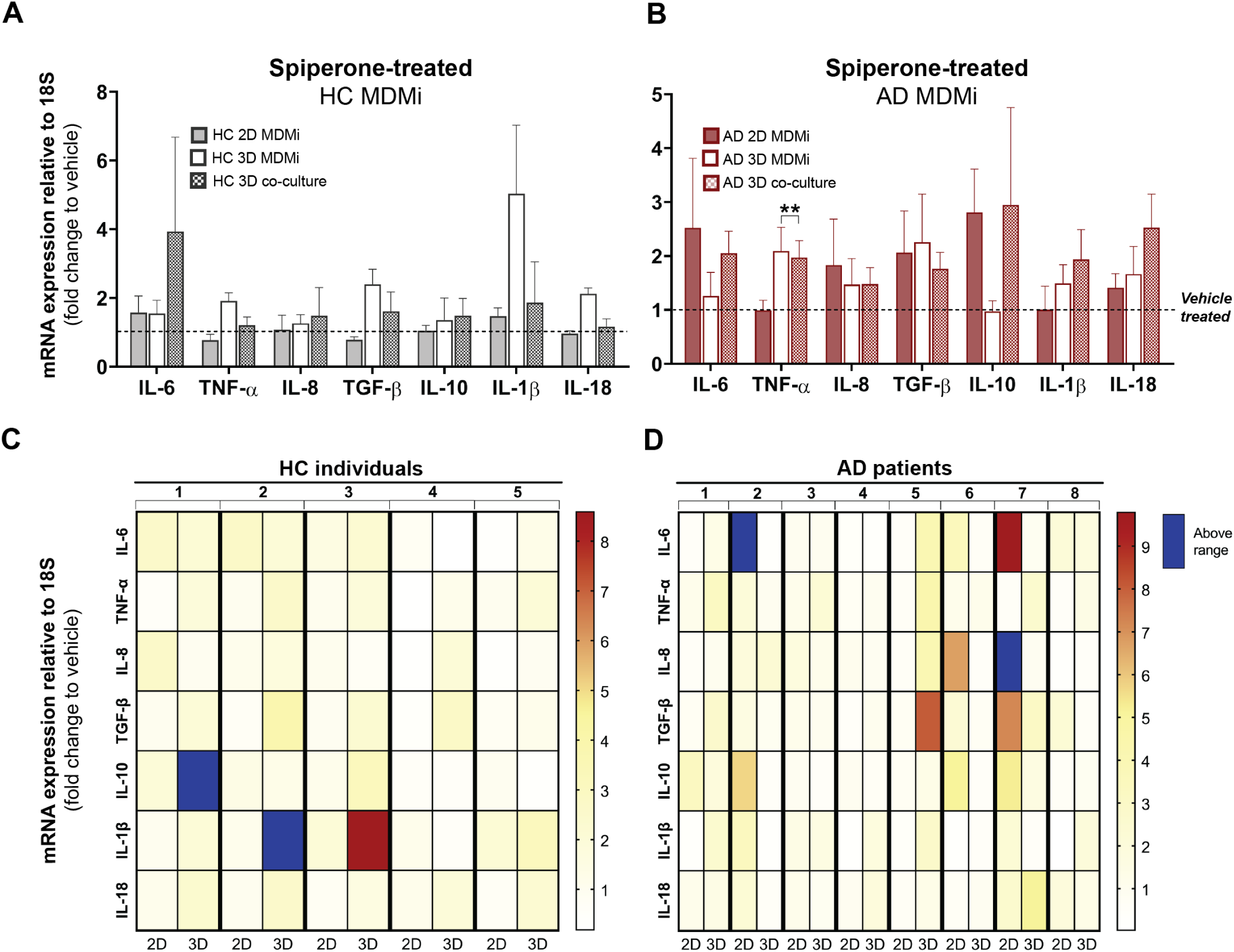
Spiperone treatment induces cytokine expression responses that differ between culture format and are heterogeneous among HC and AD MDMi. Fold change in cytokine mRNA expression levels following 24 h exposure to 1µM spiperone compared to vehicle (DMSO)-treated cultures in **(A)** HC (*n* = 5) and **(B)** AD (*n* = 8) 2D and 3D MDMi mono-cultures and 3D co-cultures. Dotted black lines represent baseline responses of vehicle-treated cultures. Heat maps showing **(C)** HC (*n* = 5) and **(D)** AD (*n* = 8) donors-specific changes in mRNA expression from MDMi mono-cultures in 2D and 3D. Red-yellow colour spectrum represents relative fold change of mRNA expression after spiperone treatment compared to vehicle. Expression changes falling outside the displayed range are indicated in dark blue. Data are presented as mean ± SEM. One-way ANOVA with Dunnett’s multiple comparison test; \*\**P* < 0.01.

